# APP accumulates around dense-core amyloid plaques with presynaptic proteins in Alzheimer’s disease brain

**DOI:** 10.1101/2020.10.16.342196

**Authors:** Tomàs Jordà-Siquier, Melina Petrel, Vladimir Kouskoff, Fabrice Cordelières, Susanne Frykman, Ulrike Müller, Christophe Mulle, Gaël Barthet

## Abstract

In Alzheimer’s disease (AD), a central role is given to the extracellular deposition of Aβ peptides, remotely produced by the proteolysis of the amyloid precursor protein (APP). This contrasts with other neurodegenerative diseases which are characterized by the intraneuronal aggregation of full-length proteins such as huntingtin, α-synuclein or TDP-43. Importantly, the distribution of APP around amyloid plaques is poorly characterized. Here, we combined an extensive set of methodological and analytical tools to investigate neuropathological features of APP in the human AD hippocampus and in two mouse models of AD. We report that APP remarkably accumulates in the surrounding of dense-core amyloid plaques together with the secretases necessary to produce Aβ peptides. In addition, the Nter domain, but not the Cter domain of APP is enriched in the core of amyloid plaques uncovering a potential pathological role of the secreted APP-Nter in dense-core plaques. To investigate the subcellular compartment in which APP accumulates, we labelled neuritic and synaptic markers and report an enrichment in presynaptic proteins (Syt1, VAMP2) and phosphorylated-Tau. Ultrastructural analysis of APP accumulations reveals abundant multivesicular bodies containing presynaptic vesicles proteins and autophagosomal built-up of APP. Altogether, our data supports a role of presynaptic APP in AD pathology and highlights APP accumulations as a potential source of Aβ and Nter peptides to fuel amyloid plaques.

## Introduction

Alzheimer’s disease (AD) is a neurodegenerative disease (ND) characterized by a severe cognitive impairment and by a prominent injury of the hippocampus. Despite important breakthroughs in the genetic and in the cellular biology of AD pathology, the etiology of the disease is still unknown and today no curative therapy is available. The brain of AD patients presents numerous amyloid plaques, hallmarks that are at the basis of the amyloid cascade hypothesis [27], a theory supported by compelling genetic evidence. Indeed, the early-onset hereditary form of the disease (FAD: familial AD) is caused by missense mutations either in the amyloid precursor protein (APP) [23] or presenilin [70], respectively the substrate and the enzyme necessary for the production of Aβ peptides which compose the amyloid plaques [13].

Remarkably, AD stands apart from the other NDs by the extracellular location of amyloid plaques which are proposed to be formed by the deposition of remotely produced Aβ peptides circulating in the extracellular fluids [18, 53]. In contrast, other NDs have in common to be proteinopathies characterized by the aggregation and accumulation of a main culprit protein inside neurons [62, 76]. For example, Parkinson’s disease, Huntington’s disease, amyotrophic lateral sclerosis and frontotemporal dementia are characterized by the aggregation of α-synuclein, Huntingtin and TDP-43 proteins within intraneuronal inclusion bodies. Moreover, the pathological mechanisms involved in AD are also proposed to involve extracellular processes. Indeed, the amyloid cascade theory states that before precipitating in plaques, Aβ peptides of different forms circulate in the extracellular space and initiate neuronal injuries by binding cell surface receptors located at synapses [67].

This central role attributed to a by-product of APP degradation, Aβ peptides, also contrasts with the mechanisms involved in the other NDs in which full-length proteins are misfolded. Yet, the presence of APP around plaques was reported soon after the cloning of APP [1, 10, 11, 71], but the relevance of this localization to AD pathology has been mainly overlooked. Notably, these early reports were mostly descriptive and their interpretation was limited by the absence of immunological controls and by the lack of quantitative analysis. Mostly based on chromogenic stainings, the early descriptions of APP around plaques lacked a detailed molecular characterization allowed by immuno-fluorescent co-labelings. Importantly, the extent and arrangement of APP accumulations in defined anatomical regions of the human hippocampus, their content in APP secretases, their molecular compositions relative to presynaptic versus post-synaptic proteins, and their ultrastructural composition have not been quantitatively investigated.

Here we combined triple-immunofluorescence labeling and electron microscopy in hippocampal sections from a cohort of 27 human patients and from the APP/PS1 and APP-NL/G/F mouse models of AD aiming to address the distribution of APP and its fragments in an unbiased approach. We report that APP, together with β- and γ-secretases, preferentially accumulate around dense-core amyloid plaques containing Aβ and APP-Nter in the hippocampus of AD patients. In addition, APP accumulations contain presynaptic markers and phosphorylated Tau in multivesicular membranes and their abundancy correlates with neuronal loss. These features support a key role of intraneuronal APP in AD pathology and suggest that APP accumulations are potential degradation sites for the formation of amyloid plaques.

## Results

### APP accumulations in the hippocampus of AD patients

We compared the distribution of APP in the human hippocampus of AD cases versus controls, by collecting paraffin-embedded hippocampal sections from two brain banks: the Netherlands brain bank (NBB) and the Karolinska Institutet brain bank (KIBB) (see Methods). Three groups were constituted: 11 controls patients (mean age 79.6 years old, 73%F and 27%M), 12 sporadic AD patients (mean age 81 years old, 92%F and 8%M) and 4 FAD patients (mean age: 48,5 years old, 50%F and 50%M) (summarized in Table 1; detailed clinical data tables in TableS1-3).

We first labeled APP using two specific antibodies against the APP C-terminal domain (APP-Cter) the C1/6.1 antibody and the Y188 antibody (Figure 1A). Preadsorption with a recombinant APP-Cter peptide abolished the staining (Figure S1A-B). In sections from the control group, APP appeared to be mostly abundant in neuronal soma using a detection with 3,3’-diaminobenzidine (DAB) as a chromogen (Figure 1B-C, S1C). In the hippocampus, the dentate gyrus granule cell layer (GCL) and the pyramidal cell layer in CA4 and CA3 are characterized by a dense DAB staining of APP (Figure 1B-C, S1C). Surprisingly, in both FAD and SAD conditions, the somatic DAB staining of APP appeared faint in the different neuronal layers revealed by the hematoxylin staining (Figure 1B, S1C). In the FAD and SAD groups, APP was prominent in remarkably intense stainings, which appeared as extra somatic protein accumulations spread in the different hippocampal regions (Figure 1B, S1C). In close-up acquisitions, in the molecular layer (ML) of the dentate gyrus or in the CA4 region, the labeled structures appear like intense clusters of APP (Figure 1C and S1D).

**Figure 1.**
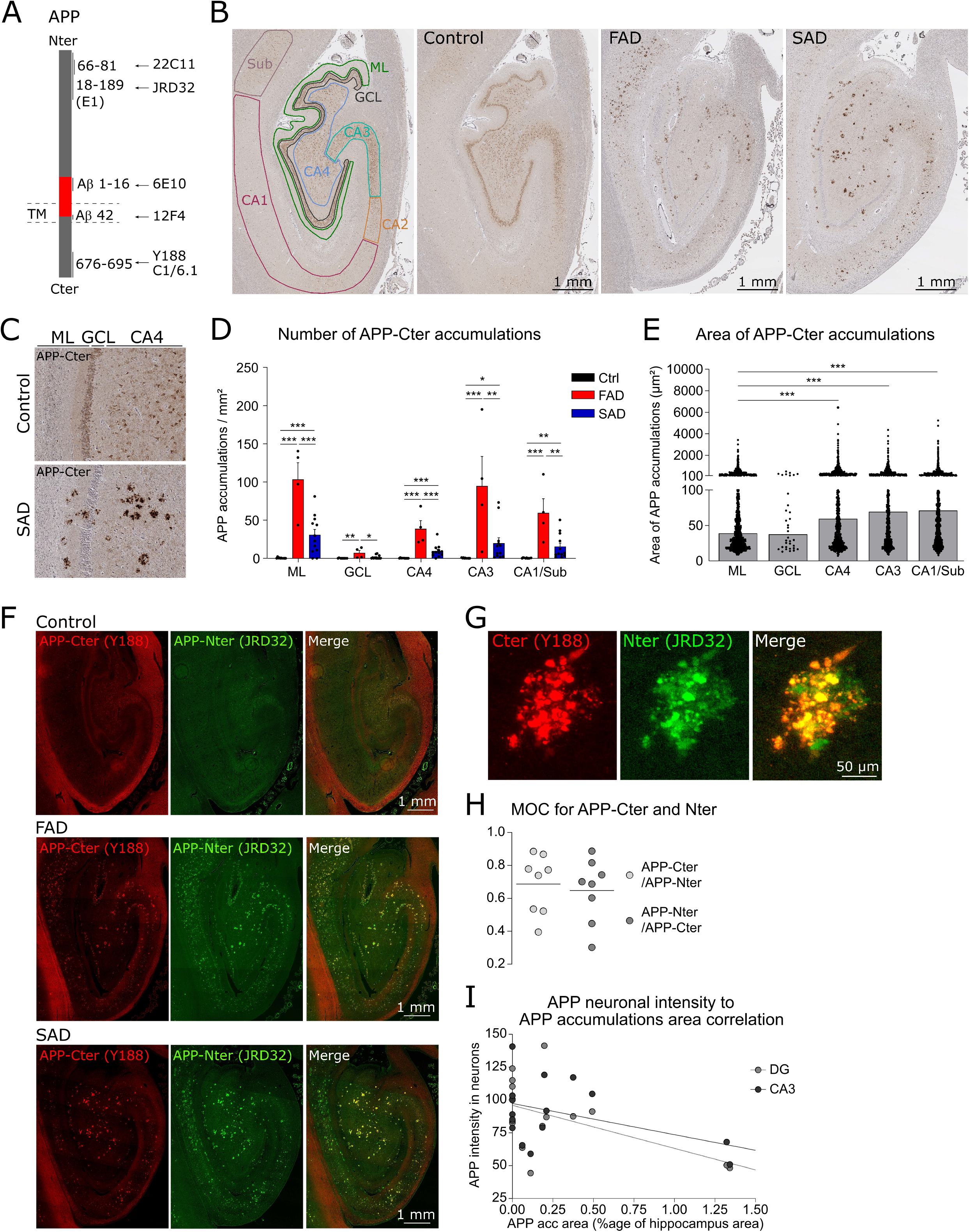
APP accumulations in the hippocampus of AD patients. (A) Scheme of full-length APP showing the epitopes in the Nter, Cter and Aβ domains of six antibodies against APP. The red segment represents the Aβ sequence spanning through the transmembrane (TM) domain. (B) Pictures of human hippocampal sections stained by DAB against APP-Cter with C1/6.1 antibody. Cases from control, FAD and SAD groups are illustrated. The first panel on the left depicts different anatomical regions where the subsequent quantifications have been performed. In controls, APP is stained mostly in the soma of neuronal layers of the dentate gyrus granule cells (GCL), CA4 and CA3. In both FAD and SAD cases, intense extra-somatic APP stainings are observable. (C) Image magnification of the dentate gyrus as in ‘B’. In the SAD case, intense extra-somatic APP accumulations are observable in the dentate gyrus molecular layer (ML), GC and CA4 regions. (D) Histogram of the number of APP-Cter accumulations in different hippocampal regions in the control (ctrl), FAD or SAD conditions. APP accumulations are barely detected in Ctrl whereas they are abundant in AD. The number of accumulations is higher in FAD compared to SAD condition. (E) Histogram of the area of APP accumulations in different hippocampal regions. The CA regions contain larger APP accumulations than the ML. (F) Human hippocampal sections fluorescently immuno-labeled against APP-Cter with Y188 antibody and APP-Nter with JRD32. In controls, both Cter and Nter APP antibodies mostly label the soma of neurons in the GCL, CA4 and CA3. In addition, Y188 nonspecifically stained the white matter tracts (see FigS1C). In both FAD and SAD cases, intense extra-somatic APP stainings, co-labeled with the two antibodies are observable. (G) Close-up picture of an APP accumulations co-labeled with APP-Cter and Nter antibodies. Scatter plot of Manders overlapping coefficients (MOC) (Method described in FigS1L-M) between APP-Cter and Nter labelings. The analysis revealed a high colocalization between the two channels suggesting the presence of full-length APP in the accumulations. (I) Graph of the mean pixel intensity of APP staining in neuronal somas in function of the area covered by APP accumulations (normalized to the hippocampus area). The linear regression of the data presents a negative slope. The Pearson correlation coefficient was −0.53 for DG (p=0.035), −0.43 for CA3 (p=0.046), indicative of an inverse correlation between the extent of APP accumulations and APP intensity in the neuronal soma.

The density of APP accumulations was considerably higher in the hippocampus of AD patients compared to hippocampus from controls, in which they were almost never detected (Figure 1D, S1E). Moreover, more APP accumulations were detected in the hippocampus of FAD patients as compared to SAD in the molecular layer (ML), CA3 and CA1 regions (Figure 1D, S1E) suggesting a relationship between the number of APP accumulation and the severity of the disease. Besides, the APP accumulations were larger in the cornus ammonis (CA) than in the ML (Figure 1E, S1F).

While giving an excellent sensitivity due to the enzymatic amplification of the detection, the DAB staining approach considerably limits the molecular characterization of the detected objects. Consequently, we developed an immuno-fluorescence approach combining the antigen retrieval method and autofluorescence quenching to perform triple labeling in human brain tissue (see Method and Controls in Figure S1G). We first verified the specificity of the two antibodies against APP-Cter by comparing fluorescence intensity in WT versus APP-KO mice. We observed that the C1/6.1 antibody was fully specific showing no labelling in APP-KO mouse brains (Figure S1H). The Y188 antibody was also specific in the grey matter without any neuronal labeling in APP-KO mouse brains (Figure S1I). However, some white matter tracts remained fluorescent in the APP-KO condition revealing that the white matter staining was not specific for APP (Figure S1I, Figure S1C). We first co-stained human hippocampal sections with both antibodies against the APP-Cter (C1/6.1 and Y188) and observed a high colocalization between the two antibodies within APP accumulations (Figure S1J, K). We developed a method, based on the subcellular colocalization analysis JACoP [8], to calculate the Mander’s overlap coefficient (MOC), an indicator of colocalization between two channels (Figure S1L, M). The proportion of pixels positive for C1/6.1 labelling which overlapped the pixels positive for Y188 was defined as MOC (C1/6.1 / Y188) and was 0.95. The proportion of Y188 pixels overlapping the C1/6.1 pixels was defined as MOC (Y188 / C1/6.1) and was 0.74 (Figure S1N). This nomenclature of MOCs was used in the entire manuscript. Then, we measured the intensity of the fluorescent labeling of APP-Cter inside versus outside accumulations. APP labeling was nine times brighter inside accumulations indicating a considerable enrichment in APP within these hallmarks (Figure S1O).

The observation of accumulations with the APP-Cter antibodies raised the question of whether the APP accumulations also contain the Nter counterpart of APP, which may suggest the presence of full-length APP. To address this question, we labeled the human hippocampal sections with two different antibodies against the Nter domain of APP (Figure 1A), the standard 22C11 antibody and the highly specific JRD32 antibody (Figure S1P). From here, our study was performed on the cohort of samples from the NBB which contains eight control cases, eight sporadic and one familial cases (hereafter combined as AD). In human hippocampal sections from controls, both antibodies against the Nter domain clearly labeled neuronal soma as did the Cter antibodies (Figure 1F and S1Q). Besides, in FAD and SAD conditions, Nter antibodies strongly stained APP accumulations detected with Cter labeling (Figure 1F, G and S1Q, R). Nter and Cter antibody labelings displayed a high MOC (Figure 1H and S1S). Whereas the joint presence of APP-Cter and Nter fragments is possible within the accumulations, the high MOC suggests that the full-length APP protein is also present in the APP accumulations. We next investigated whether there was a correlation between the burden of APP accumulations, mostly present within neuropil areas, and a seeming reduction in APP expression in neuronal soma observable in the DAB staining (Figure 1B). Taking advantage of the linearity of the fluorescent signal, we measured the mean pixel intensities of APP in neurons of the GC or CA3 layers and plotted them against the surface covered by APP accumulations in each human case. (Figure 1I). An inverse correlation was observed between APP accumulations burden in neuropil areas and APP somatic expression suggesting that APP accumulations might result from a cellular misdistribution (Figure 1I).

### APP accumulations surround dense-core amyloid plaques

We further sought to characterize the topographical organization of APP accumulations in relation to amyloid plaques. For this, we revealed amyloid plaques with three parallel labelings: an antibody against the Nter of Aβ peptides (clone 6E10 against amino-acids 1-16 of Aβ), another antibody against the free Cter of Aβ42 peptides (clone 12F4) (Figure 1A) and methoxy-X04, a fluorescent derivative of Congo red. Of important note, epitopes of the C1/6.1 and Y188 Cter antibodies are distinct from the Aβ peptide sequence (Figure 1A) and thus the detection of APP accumulations cannot be due to the detection of Aβ peptides. We tested the specificity of Aβ antibodies in WT and in APP/PS1 mice in which a human APP transgene is overexpressed. The Aβ42 and the Aβ1-16 antibodies labeled amyloid plaques in APP/PS1 mice but no staining was observed in WT mice, indicating high specificity for the human Aβ peptide (Figure S2A-B). Besides, the secondary antibody alone gave no staining (Figure S2C).

We characterized APP accumulations in human brain slices in relation to the two main types of plaques, the diffuse plaques and the dense-core plaques [52], which can be distinguished by the presence of Aβ42 and by congophilic labeling (e.g. using methoxy-X04). The amyloid plaques in the hippocampus were characterized by an intense methoxy-X04 labeling in which Aβ1-16 and notably Aβ42 staining were present (Figure 2A-C and S2D-F). These plaques present APP accumulations in their periphery (Figure 2A, B and S2H) indicating that in the hippocampus, most amyloid plaques are of the dense-core type that they are characterized by APP accumulations in their vicinity. By contrast, numerous amyloid plaques were stained for Aβ1-16 but not for Aβ42 or methoxy-X04 in the human cortex of controls (Figure S2I-K). These diffuse plaques did not present APP accumulations (Figure S2J) indicating that diffuse plaques, described as benign because they are also detected in non-demented aged humans [15, 36], are not associated with APP accumulations. Inversely, APP accumulations are specific features of the harmful, dense-core amyloid plaques.

**Figure 2.**
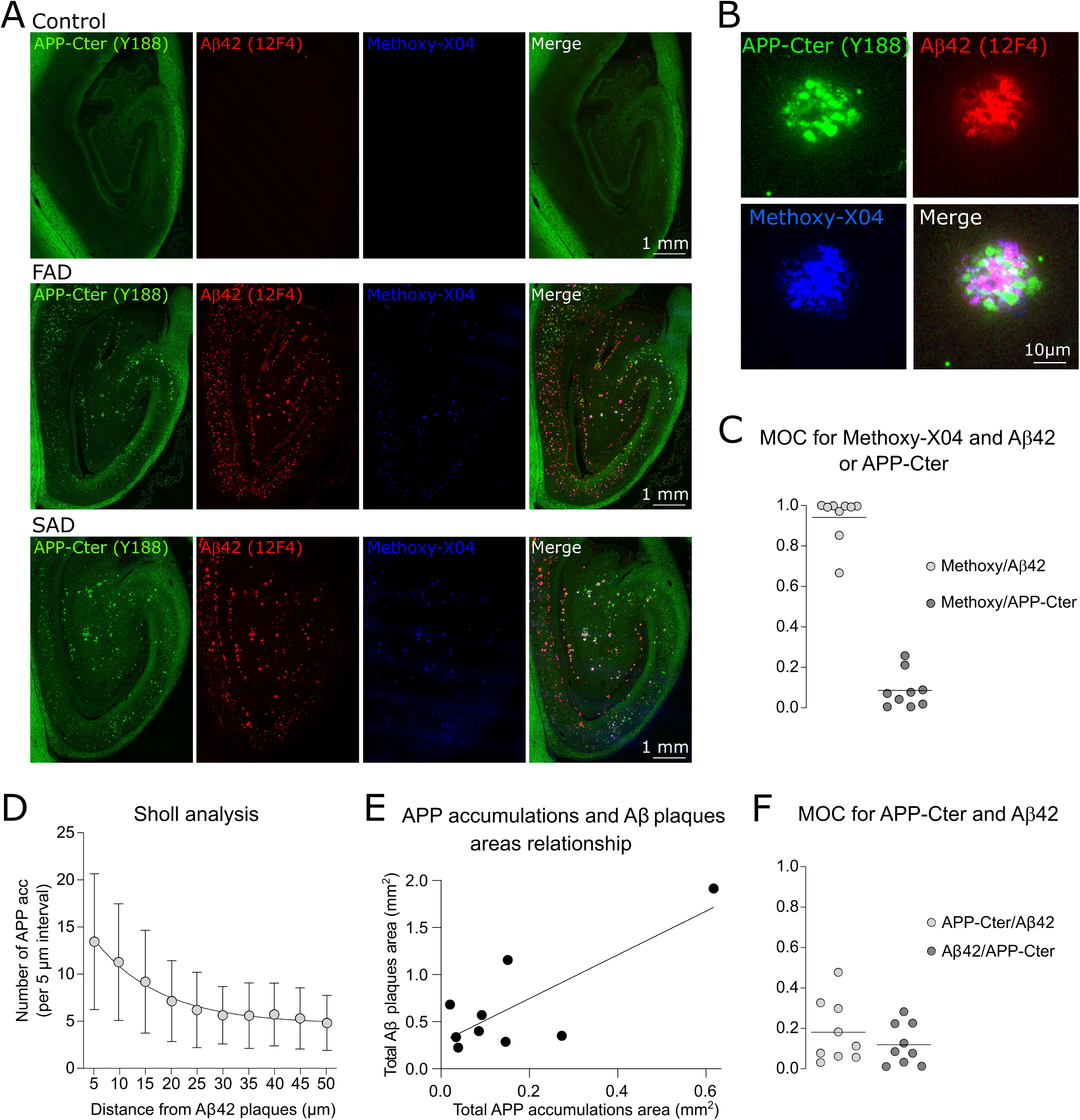
APP accumulations surround dense-core amyloid plaques. (A) Human hippocampal sections fluorescently immuno-labeled against APP-Cter and Aβ42 with respectively Y188 and 12F4 antibodies and with methoxy-X04. In both FAD and SAD cases, intense Aβ42 and methoxy-X04 labelings are observable in close proximity to APP accumulations whereas no staining is observable in control. (B) Close-up images of APP accumulations surrounding an amyloid plaque co-labeled with Aβ42 and methoxy-X04. Aβ42 and methoxy-X04 colocalize well together but not with APP-Cter. (C) Scatter plot of the MOC between methoxy-X04 and Aβ42 or APP-Cter. The analysis revealed a high colocalization between methoxy-X04 and Aβ42 but not with APP-Cter indicating that amyloid plaques and APP accumulations are located nearby but non-overlapping. (D) Graph of a Sholl analysis of the number of APP accumulations in function of the distance from the edge of Aβ42 plaques. More APP accumulations are present close to the amyloid plaques. (E) Graph of the total area covered by amyloid plaques in the hippocampus of AD cases in function of the total area of APP accumulations. The linear regression of the data presents a positive slope. The Pearson correlation coefficient was 0.79 (p=0.011), indicative of a positive correlation between the amyloid load and the extent of APP accumulations. (F) Scatter plot of MOC between APP-Cter and Aβ42. The analysis revealed a low colocalization between the two channels indicating that amyloid plaques and APP accumulations are non-overlapping.

In an effort to better assess the distribution of APP accumulations relative to amyloid plaques, we performed a Sholl analysis [72] to count the number of APP accumulations in intervals of 5 μm, up to 50 μm away from amyloid plaques used as references (Figure S2L). This analysis revealed that APP accumulations surround dense-core amyloid plaques, with a progressive decrease away from the plaque’s boundaries (Figure 2D). Interestingly, the amyloid load correlates with the extent of APP accumulations, the total area covered by amyloid plaques in the hippocampus being larger in patients exhibiting a larger total area of APP accumulations (Figure 2E). Moreover, Bland-Altman analyses of the difference of areas covered by Aβ42 and APP accumulations plotted against their average areas revealed that the data sets were of good agreement (Figure S2M). This indicates that APP accumulations and amyloid plaques probably are pathologic features which are not independent of each other even though they do not colocalize (Figure 2F).

### APP accumulations contain the secretases to produce Aβ peptides and APP-Nter enriched in their amyloid core

The detection of both APP–Cter and –Nter around dense-core plaques raises the possibility that proteolysis of full-length APP within accumulations plays a role in the generation and or maintenance of amyloid plaques. We thus characterized the distribution of the enzymatic machinery necessary to cleave APP in the accumulations. In the control hippocampus, BACE1 (β-secretase) was prominently observed in neuronal soma reminiscent of the expression of APP, in addition to a fainter labeling of the neuropil (Figure 3A). In AD condition, BACE1 was found around amyloid plaques as previously reported [34, 89] (Figure 3A, B). More precisely, it was located in APP accumulations but not inside methoxy-X04 positive fibrillary plaques, as revealed by the MOC of APP-Cter over BACE1 of 0.68 (Figure 3C) whereas the MOC of methoxy-X04 to BACE1 overlap was of only 0.05 (Figure S3A). Moreover, the average intensity of BACE1 labeling within APP accumulations was 84% higher compared to the average intensity 15 microns outside the edge of the accumulations, demonstrating the enrichment of BACE1 within the APP accumulations (Figure 3D).

**Figure 3.**
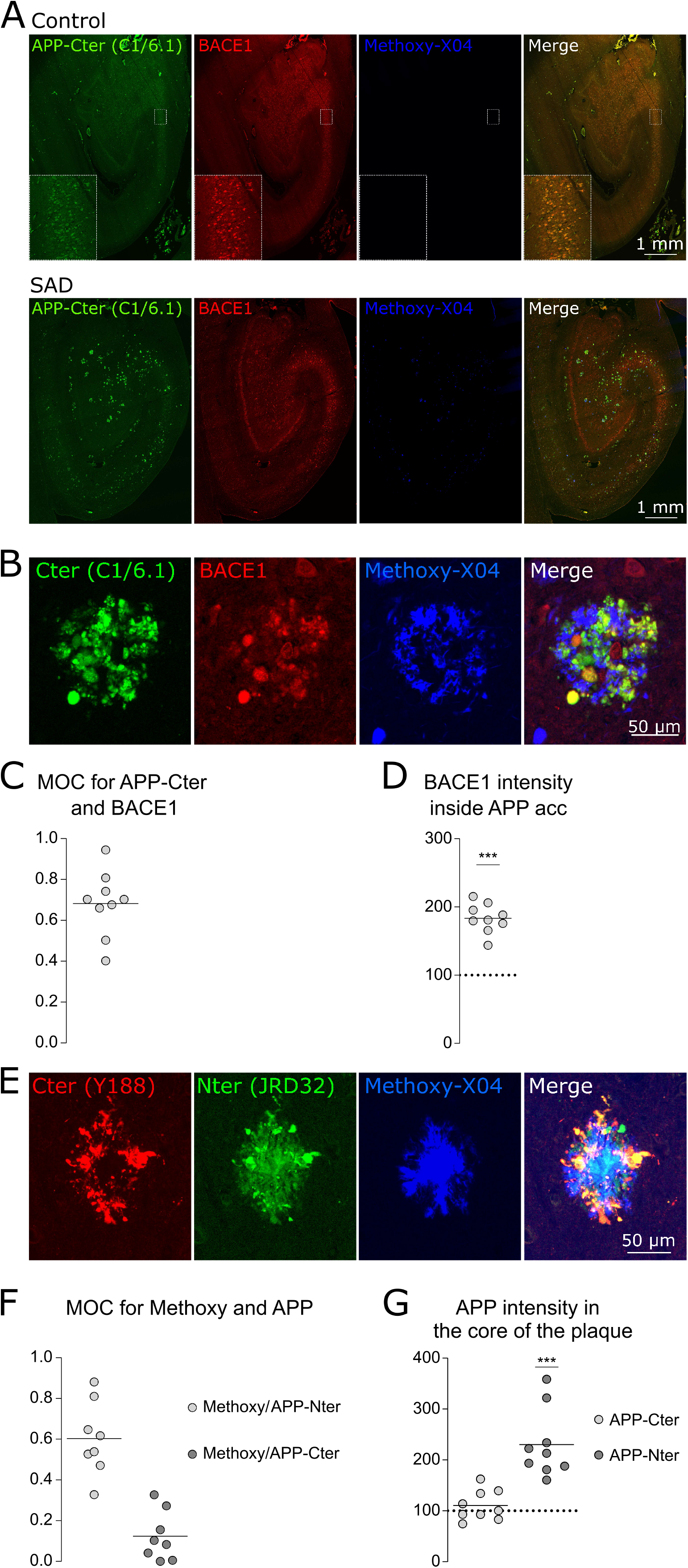
APP accumulations contain the secretases to produce Aβ peptides and APP-Nter enriched in the core of amyloid plaques. (A) Human hippocampal sections immuno-labelled against APP-Cter, BACE1 and methoxy-X04. BACE1 is prominently observed in neuronal soma reminiscent of the expression of APP, in addition to a fainter labeling of the neuropil. In both FAD and SAD cases, BACE1 staining is observable in APP accumulations. (B) Close-up pictures of stainings as in ‘A’ showing BACE1 and APP accumulations colocalizing in the surrounding of an amyloid plaque labeled with methoxy-X04. (C) Scatter plot of the MOC between APP-Cter and BACE1. (D) Scatter plot of the intensity of BACE1 labeling within APP accumulations. The data are normalized to the intensity outside the APP accumulations represented here by a dashed line. The analysis revealed that BACE1 is enriched inside the APP accumulations. (E) Close-up pictures of hippocampal amyloid plaques labelled with APP-Cter, Nter and methoxy-X04 revealed a remarkable abundance of APP-Nter in the core of dense-core amyloid plaques surrounded by APP. (F) Scatter plot of the MOC between methoxy-X04 and APP-Nter or Cter. The analysis revealed a high colocalization between methoxy-X04 and APP-Nter but not with APP-Cter. (G) Scatter plot of the intensity of APP-Cter or Nter labelings within the core of the plaques labeled with methoxy-X04. The data are normalized to the intensity outside the plaques and accumulations represented here by a dashed line. The analysis revealed that APP-Nter is enriched in the core of the plaques whereas APP-Cter is not.

The high abundancy of APP and BACE1 in the same locations suggests the possibility that APP-Nter domains (e.g. sAPPα/β) are released from the accumulations. Co-stainings for APP-Nter together with methoxy-X04, unexpectedly revealed a remarkable abundance of APP-Nter in the core of amyloid plaques (Figure 3E and S3B). The colocalization between APP-Nter and the core of the plaques was specific of the Nter domain since the MOC between methoxy-X04 and APP-Nter staining was 0.6 whereas the MOC between methoxy-X04 staining and APP-Cter was only 0.14 (Figure 3F and S3C). Moreover, the intensity of the fluorescent labeling of APP-Nter inside amyloid plaques was 2.3 times brighter than in the neuropil, indicating a marked enrichment of APP-Nter inside dense-core plaques (Figure 3G). These results suggest that in addition to Aβ, APP-Nter can accumulate at the core of the plaques possibly representing another aggregated APP derivative in addition to Aβ.

We then investigated whether γ-secretase, the second APP protease necessary to produce Aβ was also present within APP accumulations. We labeled PS1, the catalytic subunit of γ-secretase, using 33B10 antibody [21]. In all conditions, the most prominent labeling was found at the level of neuronal soma (Figure S3D). In AD conditions, PS1 staining was also found overlapping with APP accumulations (Figure S3D, E) with a moderate MOC of APP-Cter to PS1 overlap of 0.4 (Figure S3F); the MOC of methoxy-X04 to PS1 overlap was of 0.16 (Figure S3H). Moreover, comparative analysis of staining intensity revealed a modest enrichment of PS1 within APP accumulations (Figure 4H; 20% increase over the intensity outside APP accumulations). Altogether, we report that the complete machinery necessary to produce Aβ, i.e. the substrate APP and the proteolytic enzymes BACE1 and PS1, is present around plaques and could serve as local Aβ production sites.

**Figure 4.**
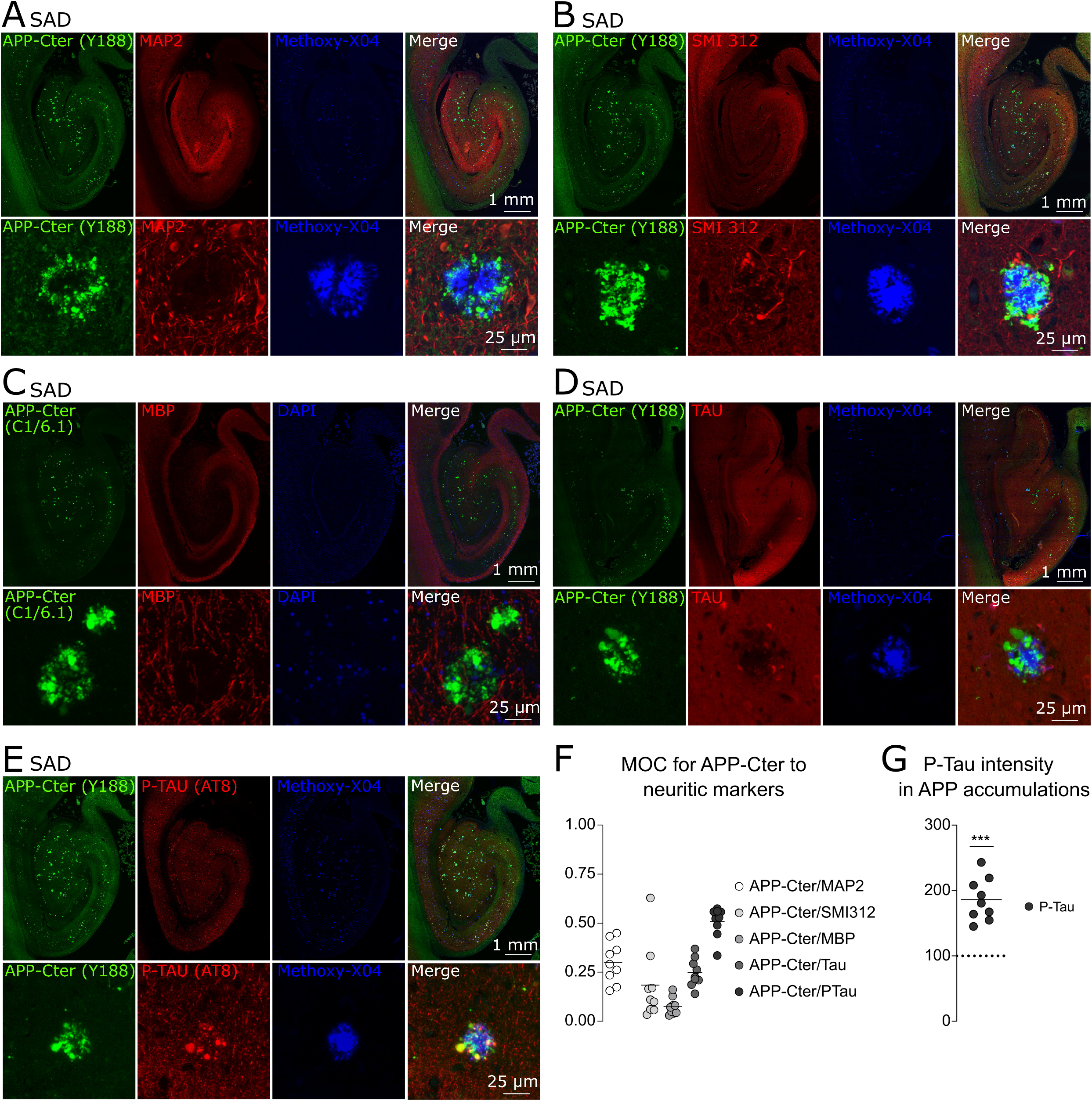
Phospho-Tau concentrate in APP accumulations. (A) Pictures of a human hippocampal section labeled against APP-Cter and MAP2 and with methoxy-X04. The lower panels show a hippocampal dense-core plaque (revealed by methoxy-X04) and surrounded by APP-Cter. It corresponds to a dark area in the MAP2 channel. (B) Pictures as in (A) but with SMI312. The lower panels show a hippocampal dense-core plaque (revealed by methoxy-X04) and surrounded by APP-Cter. Some SMI312 positive neurites surrounds the plaques but do not colocalize with APP-Cter. (C) Hippocampal sections stained for APP, MBP and DAPI. APP accumulations are mostly found out of regions labelled for MBP. The close-up pictures show the absence of colocalization between MBP and APP. (D) Hippocampal sections stained for APP, Tau and methoxy-X04. Close-up pictures show few enlarged neurites positive for Tau and negative for APP. (E) Hippocampal sections stained for APP, phosphorylated-tau (P-Tau) and methoxy-X04. Close-up pictures show several enlarged neurites positive for P-Tau which colocalize with APP. (F) Scatter plot of the MOC between APP-Cter and the neuritic markers labeled in Fig4. The colocalization is low for MAP2, SMI312, MBP, and Tau but high for P-Tau (0.53). (G) Scatter plot of the intensity of P-Tau labeling within APP accumulations. The data are normalized to the intensity outside the APP accumulations represented here by a dashed line. The analysis revealed that P-Tau is enriched inside the APP accumulations.

### Phospho-Tau and presynaptic but not postsynaptic proteins concentrate in APP accumulations

In order to explore the subcellular compartments in which APP accumulates, we investigated the co-localization of these accumulations with a set of neuritic markers. Indeed, dystrophic neurites of different molecular contents (e.g. expressing phosphorylated forms of Tau (pTau) or neurofilaments), have long been reported around amyloid plaques [49, 69, 77]. We labelled: the dendritic marker MAP2, the phospho-neurofilament (SMI312), the myelin basic protein (MBP) of the myelinated axons, the non-phosphorylated (AT8) and phosphorylated forms of Tau. The distribution of these neuritic markers in controls are shown in Figure S4. The colocalization with APP accumulations was low for all these markers, except for phospho-Tau which displayed a MOC of 0.53 (Figure 4A-F) and which was enriched almost two-fold within APP accumulations (intensity ratio of 1.86) (Figure 4G).

Since neurites could correspond to either postsynaptic or presynaptic compartments, we searched for the presence of pre- or post-synaptic proteins in APP accumulations. We immunostained the post-synaptic protein Shank 2 and APP, and in parallel we detected amyloid plaques with methoxy-X04. Whereas a prominent staining of dendrites and spines could be observed (Figure 5A), we did not detect any Shank 2 within APP accumulations or plaques (Figure 5A, B and MOCs 5G and S5C). As indicated above (Figure 4A), the dendritic protein MAP2 was absent from dense-core plaques (as previously reported [12]) and APP accumulations, further indicating that APP does not accumulate in post-synaptic compartments.

**Figure 5.**
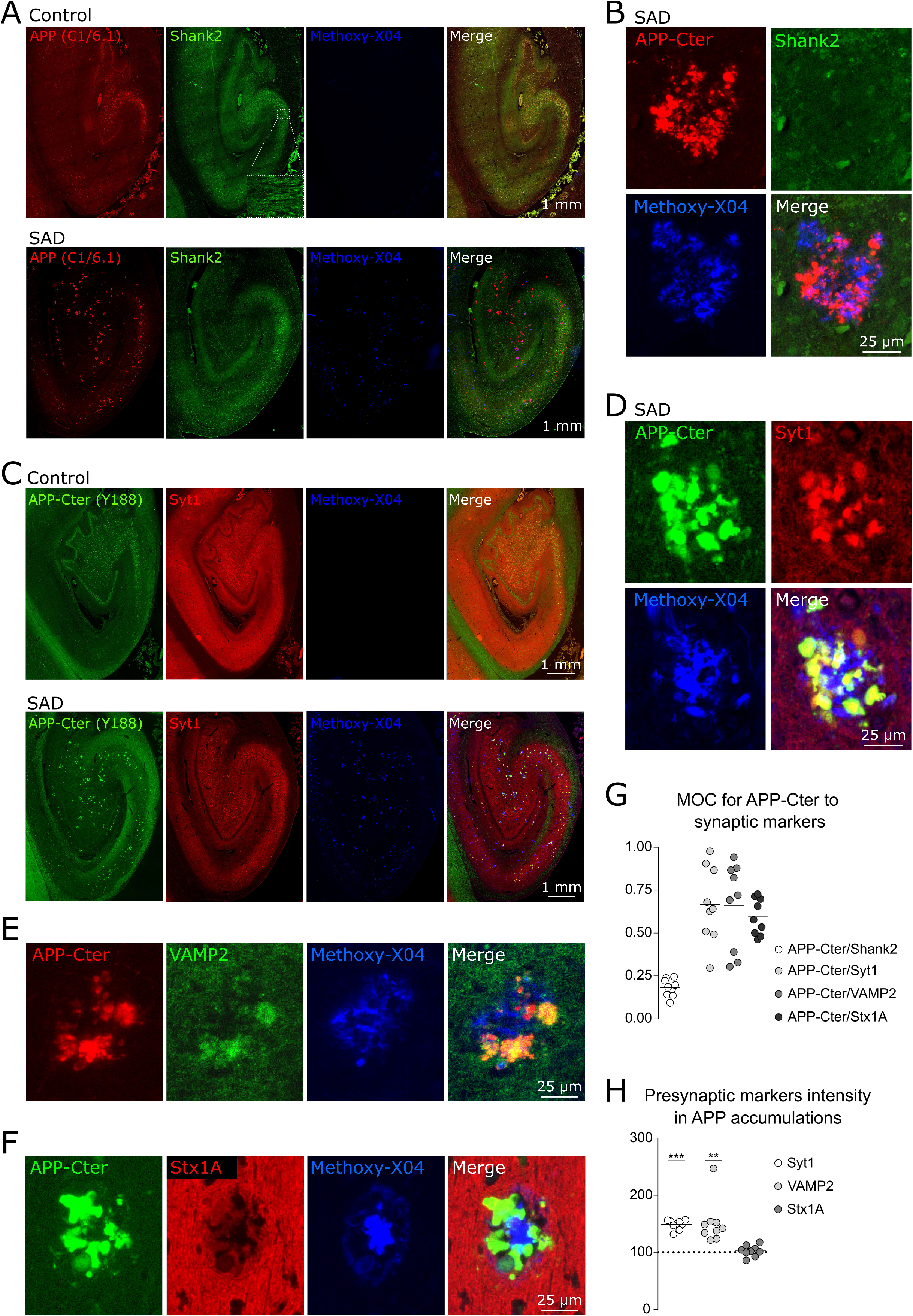
Synaptic vesicle proteins but not postsynaptic proteins concentrate within APP accumulations. (A) Human hippocampal sections immuno-labeled against APP-Cter and Shank2 and with methoxy-X04. Shank2 is prominently observed in neuronal soma and in the neuropil in both control and AD condition. (B) Close-up pictures of stainings as in ‘A’ showing the absence of Shank2 in APP accumulations surrounding an amyloid plaque labeled with methoxy-X04. (C) Human hippocampal sections immuno-labeled against APP-Cter and synaptotagmin-1 (Syt1) and with methoxy-X04. In controls, Syt1 is prominently observed in the neuropil, especially in presynaptic terminals in the CA4 region. In AD, Syt1 strikingly accumulates at the level of APP accumulations. (D) Close-up pictures of stainings as in ‘C’ showing the presence of Syt1 in APP accumulations close to an amyloid plaque labeled with methoxy-X04. (E) Close-up pictures of stainings showing the presence of VAMP2 in APP accumulations close to an amyloid plaque labeled with methoxy-X04 (full-hippocampus acquisition related to this condition are shown in FigS5A). (F) Close-up pictures of stainings showing the presence of syntaxin-1a in APP accumulations nearby an amyloid plaque labeled with methoxy-X04 (full-hippocampus acquisition related to this condition are shown in FigS5B). Scatter plot of the MOC between APP-Cter and the synaptic markers labeled in Fig5. The colocalization is high for the presynaptic markers Syt1, VAMP2 and Stx1A but low for Shank2. Scatter plot of the intensity of presynaptic markers labelings within APP accumulations. The data are normalized to the intensity outside the APP accumulations represented here by a dashed line. The analysis revealed that synaptic vesicle proteins Syt1 and VAMP2 are enriched inside the APP accumulations whereas the cell surface protein Stx1A is not.

We then stained Syt1, VAMP2 and Syntaxin1A as a set of presynaptic proteins in relation to APP accumulations and amyloid plaques. The three proteins were abundantly found in the neuropil over the entire hippocampal slice (Figure 5C, S5A, B). Interestingly, the two synaptic vesicle proteins Syt1 and VAMP2 were highly enriched within APP accumulations (Figure 5D, E, G, H) whereas they were absent from the core of the plaques (Figure S5C). Syntaxin1A, a cell-surface protein present in presynaptic active zones, was also present within APP accumulations (Figure 5F, G and S5B) although not enriched inside them (Figure 5H).

The observation that APP accumulations contain presynaptic proteins indicates that they may originate from axons or axon terminals. We questioned whether the APP accumulations were preferentially segregated in the white matter. Using MBP staining (Figure 4C and S5D) to delineate white matter tracts, we observed that APP principally accumulated in the grey matter (Figure S5E) and when found in the white matter, the accumulations were significantly smaller (Figure S5F). Collectively, these results indicate that APP accumulates together with presynaptic proteins related to vesicle release and P-Tau at presynaptic sites from unmyelinated axons.

### AD mouse models recapitulate the features of APP accumulations observed in human

In order to investigate the formation and the dynamics of APP accumulations, we aimed to study them in AD mouse models. Therefore, we first investigated whether AD mouse models faithfully recapitulated the presence of APP accumulations observed in human brains. We studied two mouse models: the APP/PS1 model which overexpresses two FAD human genes (APP with Swedish mutations and PS1ΔExon9 [32]), and the APP knock-in model NL/G/F (with four APP mutations but no overexpression [65], (Figure S6D-G)). In both models, we observed intense APP accumulations surrounding amyloid plaques, as in human (APP/PS1: Figure 6A, B and S6A-C; APP-NL/G/F: S6D, E). Besides, the APP accumulations contained the APP-Nter domain (Figure 6C, D and S6F, G; MOCs S6A) and β- and γ-secretases (Figure 6E, F) like in human AD brains. Finally, as observed in human AD brain, APP accumulations did not contain the post-synaptic protein Shank2 (Figure 6G) whereas Syt1 (Figure 6H) and Vamp2 were prominent (Figure S6H).

**Figure 6.**
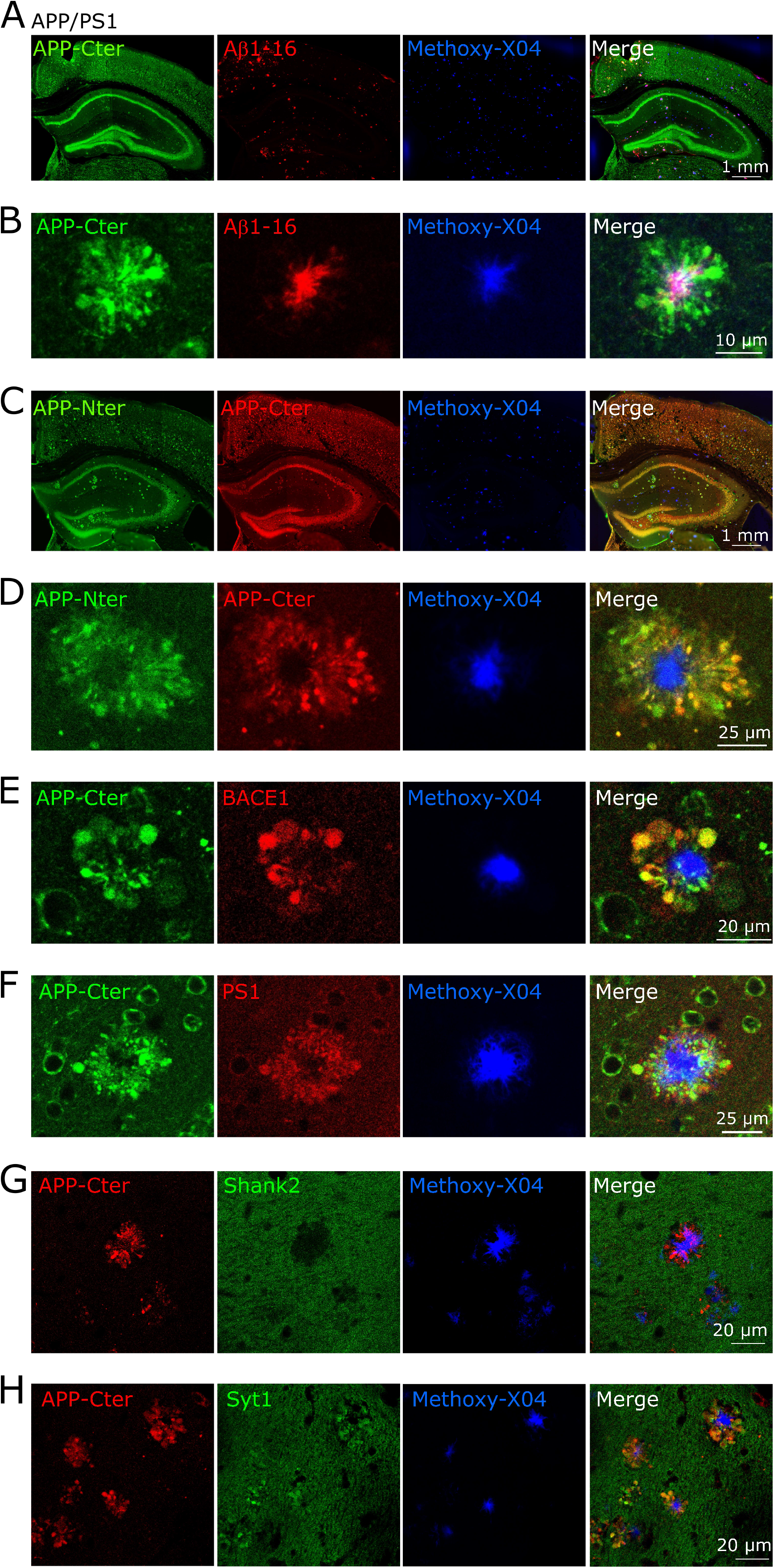
AD mouse models recapitulate the features of APP accumulations observed in human. (A, C) APP/PS1 mouse hippocampal sections immuno-labelled against APP-Cter and either Aβ1-16 in ‘A’ or against APP-Nter in ‘C’. (B, D) Close-up pictures of stainings as in ‘A, C’ showing that amyloid plaques are surrounded by APP accumulations as in human AD conditions. (E, F) Close-up pictures immuno-labelled against APP-Cter and either BACE1 in ‘E’ or PS1 in ‘F’ showing that APP accumulations contain APP-secretases necessary for Aβ production as in human AD conditions. (G, H) Close-up pictures of stainings showing the absence of Shank2 ‘and the presence of Syt1 in APP accumulations surrounding amyloid plaques labelled with methoxy-X04.

Thus, the two mouse models tested faithfully recapitulate the features of APP accumulations. From a histopathological point of view, both APP/PS1 and APP-KI NL/G/F properly exhibit APP-related AD hallmarks. However, it is noteworthy that AD-mouse models present numerous APP accumulations and dense-core plaques in the cortex, contrary to human for which the APP accumulations were more frequent of the hippocampus. Besides, the transgenic model, sometimes criticized for overexpressing the FAD genes, displayed hallmarks similar to the KI model, simply in a more abundant manner.

### Ultrastructural analyses reveal multivesicular bodies containing presynaptic proteins within APP accumulations

We further characterized APP accumulations at the ultrastructural level using electron microscopy (EM) in 9-months old female APP/PS1 mice, which exhibit abundant APP accumulations and amyloid plaques. In order to localize the structures of interest, hippocampal samples were immunostained with DAB and were then processed in osmium tetroxide and epoxy resin. We first analyzed the samples stained for APP-Nter taking advantage of its presence in both APP accumulations and plaques. We identified amyloid plaques given their distinctive shapes with radial projections at the micrometric scale [57, 84] (Figure 7A, left), and their fibrillary structures at the sub-micrometric scale (Figure 7A, right). Outside the boundaries of the amyloid plaques (Figure 7A, left), we frequently observed lysosomes connected to autophagosomes which were highly stained by the DAB reaction associated with either APP-Cter or -Nter antibody (Figure 7B). Consistent with a saturation or an impairment of lysosomal function, we observed numerous multi-vesicular bodies and autophagic vacuoles all over the tissue surrounding the plaques (Figure 7A, C), as previously reported [59].

**Figure 7.**
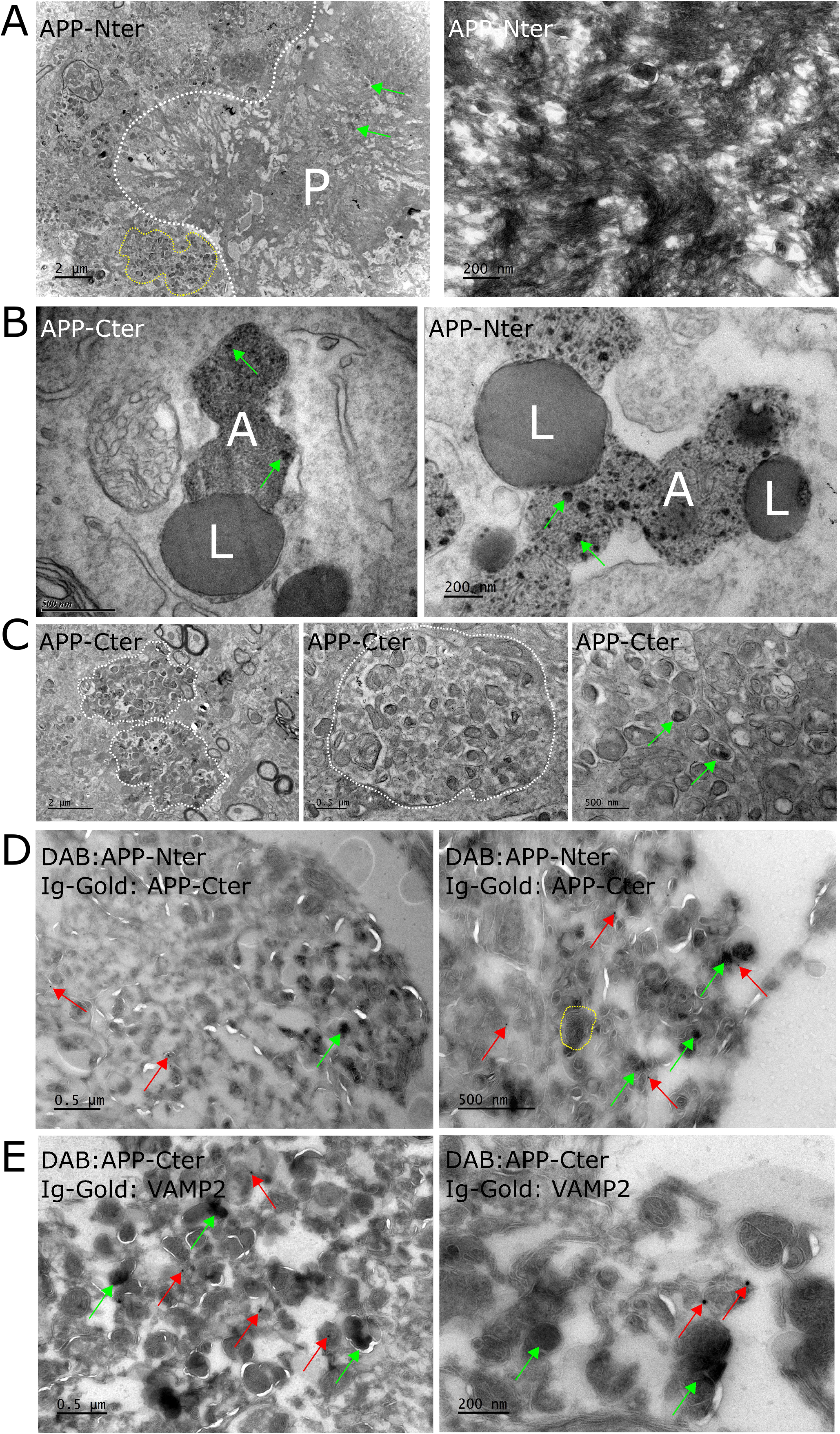
Electron microscopy analyses reveal multivesicular bodies containing presynaptic proteins within APP accumulations. (A) Electron microscopy images of a hippocampal section from an APP/PS1 mouse labeled with osmium (membrane marker) and with an antibody against APP-Nter (DAB labeling). *Left*, an amyloid plaque (P) is marked with DAB (positive for APP-Nter, green arrows) and delimited by a white broken line. Outside the plaque, round membranous organelles, positive for APP-Nter, are abundant. One cluster is highlighted inside a yellow border. *Right*, the fibrillary structure of the amyloid plaque is observable at the sub-micrometric scale. (B) Outside the boundaries of amyloid plaques, lysosomes (*L*) connected to autophagosomes (*A*) highly stained by the DAB reaction associated with either APP-Cter or -Nter antibody (green arrows) are frequently observed. (C) Clusters of multi-vesicular bodies and autophagic vacuoles are observed all over the tissue surrounding the plaques. (D) Hippocampal cryosections (Tokuyasu method) from an APP/PS1 mouse labeled with the antibody against the Nter domain JRD32 (electron dense DAB labeling, green arrows) and with the antibody against the Cter domain Y188 (antibody labeled with gold; black dots visible at high magnification, red arrows). The membranes appear finely white on the cryosections. The tissue contains numerous multivesicular bodies; one of them is highlighted inside a yellow border. They are positive for APP-Nter (DAB) and APP-Cter (Ig-Gold). (E) Hippocampal cryosections labeled with an antibody against the Cter domain of APP (DAB labeling, green arrows) and with an antibody against VAMP2 (antibody labeled with gold; black dots visible at high magnification, red arrows). The membranes appear finely white on the cryosections. The tissue contains numerous multivesicular bodies, positive for both APP-Cter (DAB) and VAMP2 (Ig-Gold).

In an effort to better characterize the membrane accumulations observed above, we performed stainings on cryosections based on the Tokuyasu method. We stained APP-Nter with DAB and labeled APP-Cter using immuno-gold particles. We observed abundant DAB staining of APP-Nter at the level of membranous accumulations (contrasted in white on dark background in cryosections) indicating that APP accumulates where membranes accumulate (Figure 7D). This observation was consistent with the presence of a transmembrane isoform of APP bearing Nter antigenicity, i.e. the full-length APP. Importantly, we observed in the same area some immuno-gold puncta revealing the presence of APP-Cter in the same location as APP-Nter (Figure 7D). This suggests again that full-length APP accumulates in membranes of the autophagy lysosomal pathway.

We then investigated the ultrastructural location of the synaptic proteins enriched within APP accumulations. Cryosections stained for APP-Cter with DAB and labeled for VAMP2 using immuno-gold particles revealed the presence of the presynaptic protein in multi-vesicular bodies positive for APP-Cter (Figure 7E). This result suggests that the multi-vesicular bodies derive from misprocessed synaptic vesicles containing APP.

### APP accumulations inversely correlate with neuronal density

The current report of APP accumulations surrounding amyloid plaques raises the question of their participation in the pathological processes in AD and first of all in neuronal loss. We first evaluated whether APP accumulations correlated with neuronal loss in the somatosensory cortex of APP/PS1. Using NeuN labeling as a neuronal reporter, we counted neurons in 100 μm-side squares, used as regions of interest (ROIs) along the sagittal to temporal axis in each slide (Figure 8A). ROIs have been placed one next to the other and did contain one, several or none APP accumulations. The analysis revealed an inverse relationship between APP accumulations and neuronal density, the larger the area covered by the APP accumulations was, the fewer neurons were present (Figure 8B). This result may be explained by neuronal dispersion due to the surface occupied by the APP accumulations within a finite area. To address this possibility, we counted the number of neurons in the full layer V from the somatosensory cortex or of the full ML of the dentate gyrus in slices from 2, 5 and 9 months-old APP/PS1 mice (Figure 8C). In both brain regions, we observed a significant and progressive decrease in the number of neurons starting at 5 months in APP/PS1 mice compared to WT (Figure 8D). This result indicated that the decrease in neuronal density observed close to APP accumulations was not caused by neuronal dispersion but by neuronal loss, in which APP accumulations may participate. Together with the observation that FAD cases exhibit more APP accumulations than SAD cases (Figure 1D), these results suggest a relationship between the number of APP accumulations and the severity of the disease.

**Figure 8.**
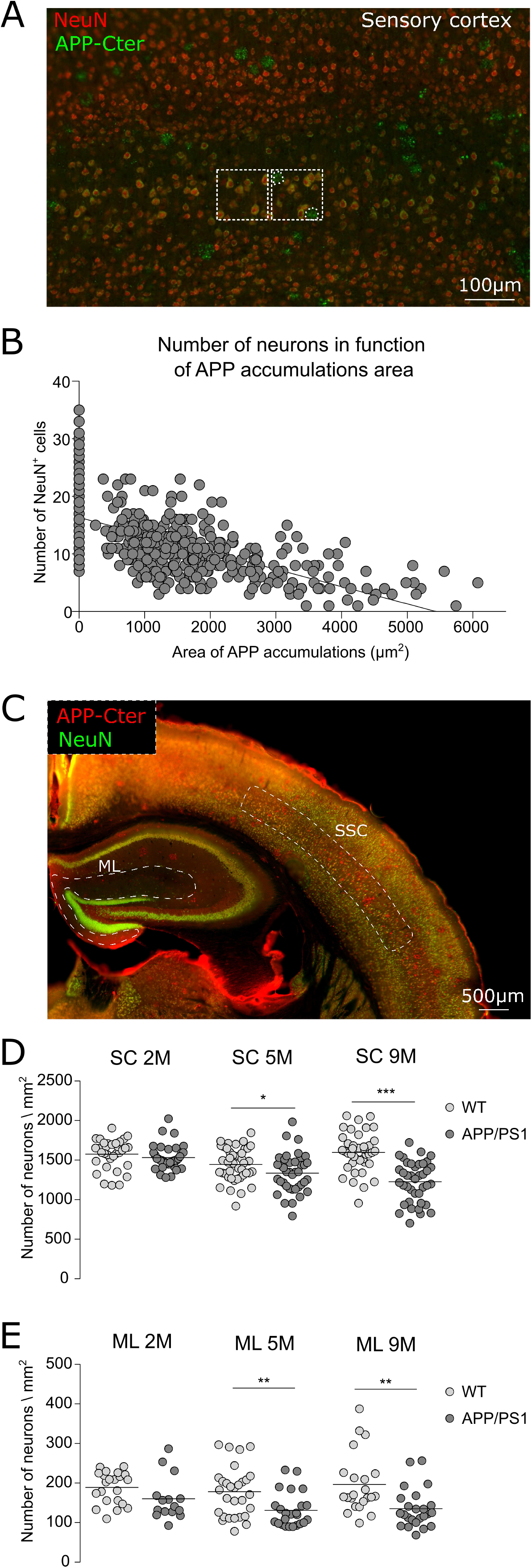
The number of APP accumulations inversely correlate with neuronal density. (A) Sections of the somatosensory cortex of APP/PS1 immuno-labeled with NeuN and APP-Cter. Using NeuN labelling as a neuronal reporter, we counted neurons in 100 μm-side squares, used as regions of interest (ROIs). The squares were placed in different regions along the sagittal to temporal axis containing or not one or several APP accumulations. (B) Analyses of pictures as in ‘A’ revealed an inverse relationship between the area covered by APP accumulations and neuronal density within the ROIs, the larger the area covered by the APP accumulations, the fewer neurons were present. (C) Pictures of APP/PS1 brain sections immuno-labeled with NeuN and APP-Cter. ROIs of the whole hippocampal molecular layer (ML) area and of the somatosensory cortex (SSC) where numbers of neurons were counted (results in ‘D’ and ‘E’) are highlighted by dashed lines. (D, E) Analyses of pictures as in ‘C’ revealed a neuronal loss in APP/PS1 at both 5 and 9 months-old in SC and in ML. The WT littermates do not present any neuronal loss.

## Discussion

The deposition of Aβ peptides within the amyloid plaques is a well-characterized feature of AD pathology. However, a large body of genetic evidence also supports the idea that the excess of other APP processing products, or FL-APP itself, is toxic for neurons in AD. For example, PS1 FAD mutations impair the proteolytic function of PS [7, 9, 78, 85] leading to the accumulation of uncleaved substrates potentially toxic for the cell [2, 5, 16, 33]. Down syndrome patients, who have three copies of the APP gene [22, 44, 47], as well as patients with a rare APP gene duplication [63, 74] suffer from early forms of AD. Finally, ApoE4, the main genetic risk factor of AD, markedly promotes APP expression [31]. However, this array of genetic evidence has been systematically used in favor of an excessive production of Aβ peptides, but not of a direct role of APP. Here by using an innovative set of tools and analyses in human brain, we performed an exhaustive qualitative and quantitative analysis of intense APP clusters which surround dense-core amyloid plaques. These APP accumulations contain presynaptic proteins and APP-secretases, and likely result from a misdistribution of proteins from neuronal soma to unmyelinated axons or terminals. These data support the idea that APP accumulations represent a potential source of the Aβ peptides composing the amyloid plaques.

### APP accumulates in brains from AD patients

The APP accumulations are detected with four different antibodies against the Cter and the Nter, which colocalize at both the micrometric and nanometric scale, suggesting that they contain FL-APP (Figure 1). The number of APP accumulations correlate with the severity of the disease, as indicated by higher incidence in FAD as compared to SAD cases (Figure 1D) and the inverse correlation between the number of FL-APP accumulations and neuronal loss (Figure 8). The molecular mechanisms by which APP accumulations may be pathogenic have not been investigated here, but it is tempting to speculate on a role of APP aggregation, i.e. excessive APP self-assembly. Although the accumulation of APP does not necessarily imply its aggregation, it is important to note that FL-APP readily dimerizes [6, 50, 56, 75]. Protein aggregation of full-length proteins (α-synuclein, Huntingtin, TDP-43) is a common feature of proteinopathies in NDs [62, 76] and it is tempting to speculate that AD proteinopathy also involves the intraneuronal aggregation of a FL-protein like other NDs in addition to the aggregation of Aβ peptide.

### APP accumulations occur within presynaptic compartments

A striking features of APP accumulations is their selective enrichment in proteins of the presynaptic vesicle machinery (Syt1 and Vamp2) (Figures 5–7), whereas uniquely postsynaptic or dendritic proteins (such as Shank2 and MAP2) appear to be absent. Recent reports indicate the accumulation of presynaptic proteins in amyloid plaques or within surrounding dystrophic neurites [26, 34]. In contrast, amyloid plaques are mostly devoid of post-synaptic proteins [26, 64]. APP shows remarkable abundance in presynaptic compartments [41, 45, 83], in which it interacts with several proteins of the synaptic release machinery [37]. This raises the possibility that APP is directly involved in the enrichment of presynaptic proteins within APP accumulations as a result of protein-protein interaction. Interestingly, the ultrastructural analysis of APP accumulations did not provide evidence for synaptic vesicles which could harbor the enriched presynaptic proteins (Figure 7), but revealed synaptic vesicle proteins residing within the numerous multivesicular bodies observed in APP accumulations.

Our observation highlights the selective accumulation of presynaptic proteins in discrete microscopic regions such as amyloid plaques or APP accumulations, even though recent discoveries show global downregulation of presynaptic proteins in AD brains [3, 28, 82]. Interestingly, we report that APP itself follows a complex pattern of misdistribution. Indeed, in addition of the accumulations found mostly within neuropil areas, we observed in AD a marked reduction in the somatic staining of APP in neuronal soma (Figure 1D, C) where APP is normally prominent. Importantly, this decrease correlates with the extent of APP accumulations burden (Figure 1I) suggesting that the two phenomena depend on the same cellular mechanisms, e.g. a slow-down of axonal retrograde transport. Indeed, an impairment of axonal transport represents a simple and straight-forward explanation for these complex variations in proteins abundance. A traffic jam would lead, on one hand, to the pathological accumulation of proteins at the site of blockage, and on the other hand, to their deficit where they exert their physiological function. Clinical and neuropathological studies have long indicated that synaptic impairment correlates strongly with cognitive deficits in AD [14, 66, 67, 79] whereas reports showed abundance in proteins involved in presynaptic plasticity like complexin 1, were associated with better cognitive function [29, 87]. Moreover, recent evidence points to a specific impairment of presynaptic proteostasis in AD [4]. How the misdistribution of proteins crucial for neurotransmitter release contributes to the impairment of memory and cognition in AD is a key question that remains to be addressed.

The presence of multivesicular bodies harboring presynaptic vesicle proteins in APP accumulations, may represent the endpoint of an axonal degeneration process. Indeed, APP accumulation is a marker of axonal damage in several pathologies including multiple sclerosis [19, 39], myelopathy [80], herpes simplex encephalitis [54], or even traumatic brain injury [20, 61]. Given the well-established fast axonal transport of APP [25, 38, 42, 73], APP may accumulate in AD as a result of axonal injury. The cause of such axonal injury could be diverse and possibly related to the destabilization of axonal microtubules by alterations (e.g. over-phosphorylation) of the microtubule-stabilizing protein Tau [81]. Indeed, phosphorylated–Tau (P-Tau) is enriched within APP accumulations but not its non-phosphorylated form (Figure 4). This observation offers the possibility of a functional interaction in the same compartment between the two main pathological actors of AD, APP and Tau, This hypothesis is supported by recent evidence that deletion of Tau reduces BACE1 around plaques and slows plaque formation in an AD mouse model [60]. In addition, P-Tau could play a pathological role related to its recently reported impairment of synaptic transmission through interaction with the synaptic vesicle synaptogyrin-3 [51, 90]. Together, our results support a role of presynaptic APP in AD pathology, within P-Tau positive dystrophic neurites, where presynaptic proteins are misdistributed, in keeping with a presynaptic failure hypothesis [4].

### APP accumulations as local sources for Aβ production

The classical paradigm of amyloid plaque formation proposes that Aβ peptides, ubiquitously produced in the brain and in the periphery, would circulate in extracellular fluids before seeding and aggregating within amyloid plaques [18, 53]. However, this rationale seems inconsistent with the long observed decrease in Aβ concentration in cerebrospinal fluid and plasma of AD patients compared to non-demented controls [55] and with the decreased proteolytic efficacy and processivity of γ-secretase containing PS1 with FAD mutations [7, 9, 78, 85].

An alternative paradigm of amyloid plaque formation, formulated by Fisher in 1907 in his early description of AD pathology, proposes that amyloid plaques derive from dystrophic, degenerating neuronal processes (reviewed in [24]). The collection of data we are presenting here, namely the striking accumulations of APP (Figure 1), their localization around dense-core plaques (Figure 2, 6), their content in secretases (Figure 3, 6), advocates that APP accumulations represent a local source for Aβ peptides composing the plaques. Indeed, the localization in the same compartments of APP and its secretases (BACE1 and PS1; Figure 3) could favor enzyme-substrate collisions and therefore Aβ production. This alternative formation of Aβ peptides is also supported by the remarkable correlation between APP accumulations and amyloid plaques (Figure 2). This model may also help understanding how abundant amyloid plaques can be formed in AD brains, despite the lower concentration of Aβ in extracellular fluids compared to controls.

### A role of APP in the accumulation of autophagic vesicles?

Multivesicular bodies (MVB) and autophagic vacuoles have previously been reported in AD [59]. Their accumulation is consistent with an impairment of the autophagy-lysosomal pathway currently debated in the AD field [40, 43, 58, 88]. Here we report the presence of APP in MVB and autophagic vacuoles (Figures 7) suggesting that APP accumulations are related to a defective degradation of APP by the autophagy-lysosomal pathway. Interestingly, several recent studies have reported the accumulation of membranes and autophagy-related organelles in NDs. For example, autophagosomes accumulate inside neurons in Huntington’s disease [48], and Lewy bodies, the main hallmarks in Parkinson’s disease, have been reported to be composed of crowded organelles and vesicles [68]. Thus, in addition to being proteinopathies characterized by the misfolding and the aggregation of proteins, NDs may be characterized by the impairment of the autophagy-lysosomal pathway. The resolution of the cause-consequence relationship between protein aggregation and lysosomal dysfunction will be one of the next challenges for research on NDs.

### APP-Nter domain as a potential therapeutic target

Despite numerous clinical trials targeting Aβ peptide (β-secretase or γ-secretase inhibitors, active or passive immunotherapy against Aβ peptides), no treatment to cure or even to slow down AD progression is available [30, 46]. There is thus, a huge need for alternative approaches in AD clinical research.

Based on the present results, we propose the Nter domain of APP as a therapeutic target of important potential for the following reasons. First, the Nter domain as part of FL-APP could play a role in the formation of APP accumulations around plaques. Second, the Nter domain is concentrated at the core of the plaques, uncovering a potential pathological role of the secreted APP-Nter in dense-core plaques. Third, because APP is first cleaved by BACE1 in the amyloidogenic pathway and then by γ-secretase, the Nter could be the first extracellular fragment, before Aβ, to be released and aggregated. Fourth, the Nter domain, due to its extracellular topology when FL-APP is located at the cell surface, is very accessible to the binding of therapeutic molecules. Such molecules could be antibodies such as the ones used in this study, or peptides, directed for example against the amino-acid sequences involved in the dimerization of APP [6], in an attempt to prevent its accumulation and aggregation.

## Acknowledgments

We thank Takashi Saito and Takaomi C. Saido from RIKEN Brain Institute (Saitama, Japan) for the gift of APP-NL/G/F brain slices. The *postmortem* human brain tissue for this study was supplied by the Netherlands Brain Bank (Amsterdam, the Netherlands) and the Karolinska Institute Brain Bank (Stockholm, Sweden); we thank all the donors for the tissue used in this study. We thank Caroline Graff from the Karolinska Institute Brain Bank. We thank Juan Pita-Almenar from Janssen Pharmaceutica for the gift of JRD32 antibody. Imaging was performed at the Bordeaux Imaging Center, a service unit of the CNRS-INSERM and Bordeaux University, member of the national infrastructure France BioImaging (ANR-10-INBS-04). This project was supported by the CNRS, it has received funding from the European Union’s Horizon 2020 research and innovation programme under grant agreement No. 676144, from the foundation Plan Alzheimer, from France Alzheimer, and from the Fondation pour la Recherche Médicale (project #DEQ20160334900).

## Author contributions

T.J-S., M.P., V.K. conducted the experiments, F.C. developed ImageJ macros for images analyses, S.F. supervised the experiments performed at the Karolinska Institute, U.M. provided research materials and advices, T.J-S., G.B. designed the experiments, T.J-S., C.M., G.B. wrote the manuscript.

## Data availability

The data that support the findings of this study are available from the corresponding author upon reasonable request.

## Methods

### Human brain tissue

The use of human brain material was approved by the ethical and research committees after the consent of patients or relatives for research use. Paraffin-embedded samples were obtained from The Netherlands Brain Bank, Netherlands lnstitute for Neuroscience, Amsterdam (www.brainbank.nl) or from and the Karolinska Institutet brain bank (KIBB). All Material has been collected from donors for or from whom a written informed consent for a brain autopsy and the use of the material and clinical information for research purposes had been obtained by the NBB.

We used sections from the hippocampus of 11 control patients (mean age 79.6 ± 6.93, 73%F and 27%M, Table 1), 12 sporadic AD patients (mean age 81 ± 9.6, 92%F and 8%M, Table 2) and 4 familiar AD cases (all cases with different PSEN1 mutations, mean age 48.5 ± 9.7, 50%F and 50%M Table 3).

### Mice

In agreement with the Bordeaux/CNRS Animal Care committee, heterozygous APP/PS1 female mice co-expressing human APP Swedish mutation (KM670/671NL) and PS1 Exon9 were used [32]. Mice were obtained from crossing heterozygous female transgenic mice with B6C3F1/J male mice. Animals were caged in groups of 4 to 10 with access to water and food. Rooms were kept at a stable temperature with 12 h light/dark cycle under pathogen-free conditions until surgery. After surgery mice were kept in single cages. Brain sections (5 μm) were also obtained from from 12-month-old female APP-KI NL-G-F mice expressing the Swedish (KM670/671NL), Iberian (I716F) and Artic (E693G) mutations [65]. Sections were fixed in PFA 4 % and mounted with paraffin.

### List of antibodies

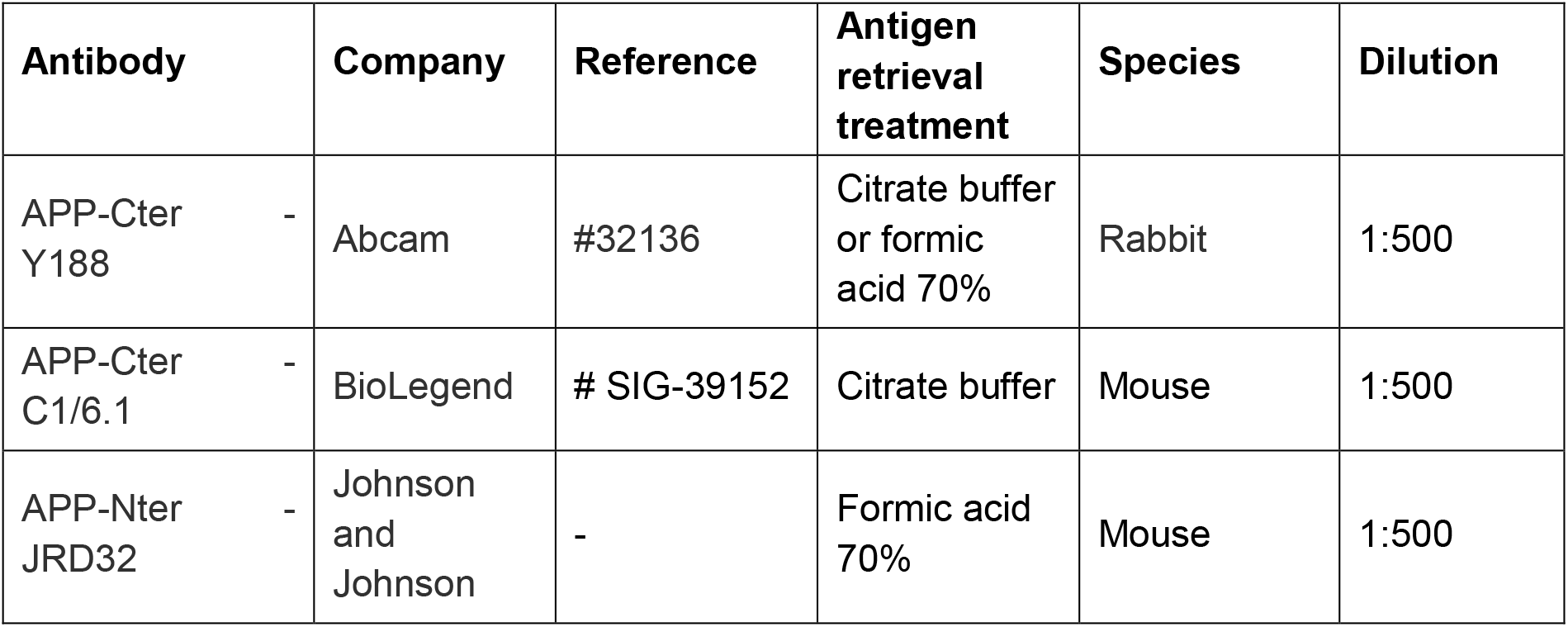
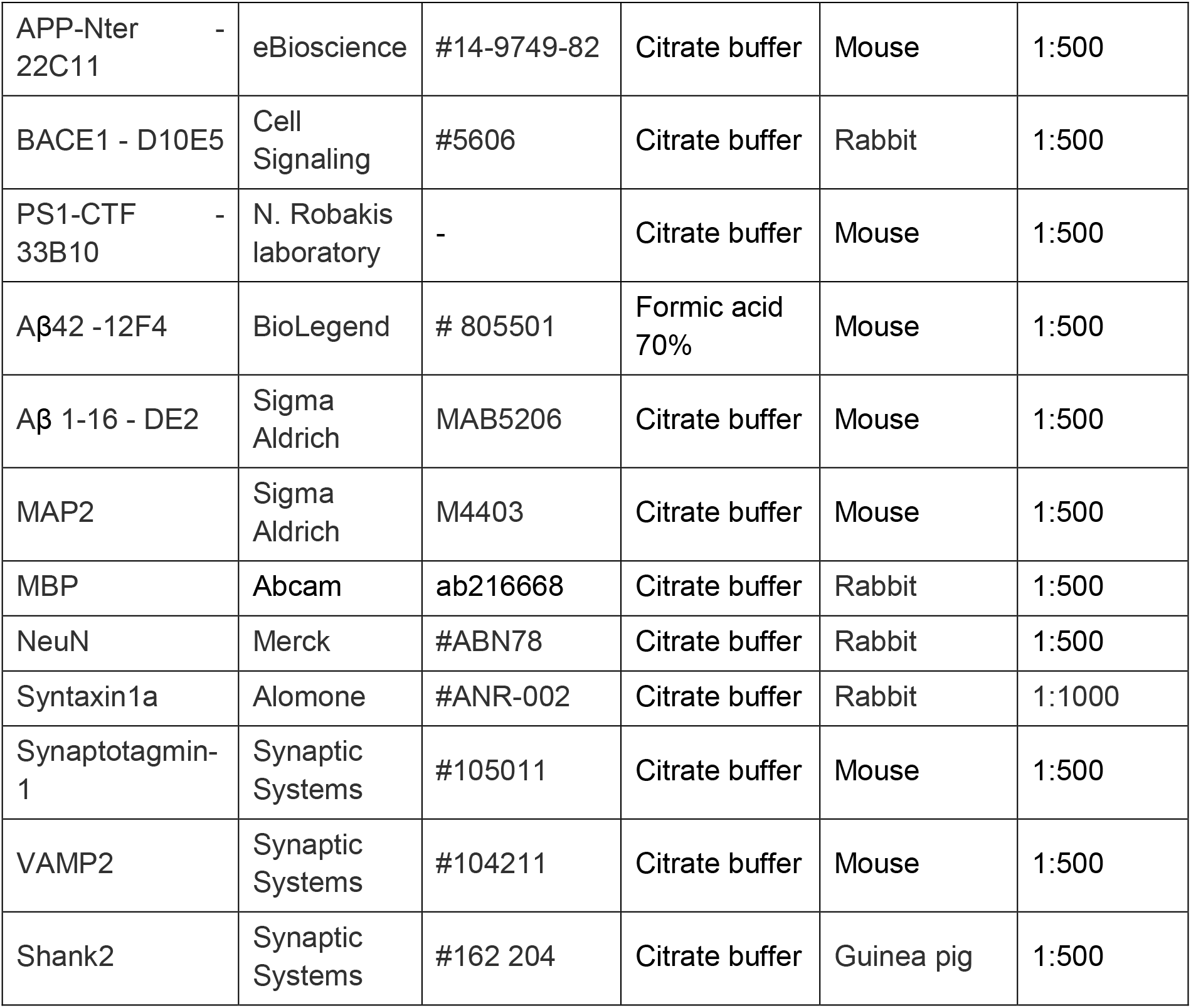

### Immunohistochemistry on human postmortem brain sections

#### DAB immunostaining

A Dako EnVision + Dual Link System-HRP (#K4065) was used to perform stainings with DAB. Paraffin-embedded hippocampus sections (5 μm) were sliced and mounted on microscope glasses. Sections were incubated 2 times in xylene (≥98.5 %, AnalaR NORMAPUR^®^ ACS, Reag. Ph. Eur. analytical reagent) for 10 minutes and hydrated for 3 minutes in different baths ranging from 99% to 70% ethanol concentrations. Then, sections were incubated in a pressure cooker (Biocare medical) at 110 °C for 30 min either in citrate buffer (10 mM Citric acid (Sigma Aldrich 251275), 0,05 % Tween-20 (Sigma Aldrich P7949), pH 6) or formic acid 70 % ((Sigma Aldrich F0507) pH 2) depending on the protein staining. Brain sections were washed with PBS-Tween 0,05 % (PBS-T) and blocked with a peroxidase enzyme blocker for 5 minutes (Dako Dual Endogenous Enzyme Block S2003) and goat serum (NGS; Thermo Fisher Scientific PCN5000) for 20 minutes at RT. Sections were then incubated at room temperature (RT) for two hours with one primary antibody. After 3 washes in PBS-Tween, sections were incubated with peroxidase-conjugated secondary antibody. Sections were then washed and incubated 2 minutes with 3,3’-diaminobenzidine chromogen solution. Sections were washed again and incubated with Hematoxylin (Sigma Aldrich #51275) for 5 min at RT before being dehydrated in baths of 3 min ranging from 70% to 99% ethanol and finally in xylene for 10 min. Sections were finally mounted with Vecta mount (VectaMount^®^ Permanent Mounting Medium #H-5000) and dried overnight. Preabsorption controls in human sections were performed by using the APP C-terminal peptide sequence KMQQNGYENPTYKFFEQMQN (Proteogenix) at 1 μg/ml and Y188 or C1/6.1 at 1 μg/ml (equal mass concentration, i.e. excess molarity of the adsorption peptide).

#### Immunofluorescence

Brain sections were sliced, hydrated and treated for epitope retrieval as for the DAB immunostaining. In addition, slices were washed and blocked with NGS, before incubation for 2 hours at RT with one or a with a combination of primary antibodies. Then, sections were washed in PBS-Tween and incubated with an autofluorescence quencher (TrueBlack^®^ Lipofuscin Autofluorescence Quencher - Biotium #23007) for 5 minutes at RT. After 3 washes in PBS 1X, sections were incubated at RT for 2 hours with secondary antibodies diluted at 1/500 (coupled with Alexa 488 or 568 - Thermo Fisher Scientific) and with Methoxy X04 (5 μM) (Tocris Bioscience #4920) or DAPI (300 nM) for 20 minutes. Finally, sections were mounted in FluoromountTM (Southern Biotech #0100-01) and dried overnight.

### Mouse immunohistochemistry

Mice were first anesthetized by an intraperitoneal injection of pentobarbital (50 mg/kg body weight). Mice were then perfused intracardially with 0.9 % NaCl for 1 min and then with 4 % paraformaldehyde in 0.1 mM PBS (PFA 4 %) for 4 min. Brains were then dissected and post-fixed for 8 h in PFA 4 %. Coronal sections of 50 μm were cut on a vibratome (Leica VT1200S) and permeabilized in PBS 0.3 % Triton X-100 (Sigma Aldrich - #9002-93-1) at RT for 2 hours. Brain sections were then incubated overnight at 4°C with the primary antibody diluted in PBS-T. In addition, sections were also incubated at RT for 5 min with an autofluorescence quencher (True black). After 3 washes in PBS 1X, sections were incubated at RT with secondary antibodies for 2 hours. Finally, slices were incubated in Methoxy-X04 or in DAPI for 10 min before getting mounted in FluoromountTM.

### Brain section imaging

Brain sections were all imaged using the bright field microscope Nanozoomer 2.0HT (Hamamatsu) with a 20X objective and fixed additional lens 1.75X. For every experiment, imaging settings and acquisitions were maintained constant between different patient groups.

### Image analysis

Image acquisitions were exported as .tiff files in independent channels by using the NDP.view2 software (Hamamatsu). Using Image J, images were then calibrated and processed as 8-bits grayscale images (0-255). A high manual threshold was first applied to the APP accumulations and plaques to remove any signal coming from other non-interesting structures. The structures of interest were then saved separately as regions of interest (ROIs) and enlarged by delineating new boundaries 15 μm away from their detection perimeter for posterior analysis. The two ROIs were then merged and used as an overlay to extract the objects from the initial image. Finally, from the extracted images, colocalization studies were performed between different channels by using the Mander’s overlap coefficient (MOC) (Figure S1J, K). Images were analyzed by using the JACop plugin (http://www3.interscience.wiley.com/cgi-bin/fulltext/118727584/PDFSTART; 2006), which provides the proportion of co-localized pixels ranging from 0 (no co-localization) to 1 (perfect co-localization). For some markers, the threshold setting was done automatically by using the plugin ‘Auto threshold’, which provides a set of different automatized algorithms. APP C-terminal accumulations and plaques (methoxy X04) were selected by using the ‘Max entropy’ algorithm that implements the Kapur-Sahoo-Wong thresholding method [35]. For Aβ antibody stainings, the algorithm ‘Yen’ was applied [86]. For Syntaxin1a and Shank2 the ‘Moments’ algorithm and for myelin basic protein the ‘Default’ algorithm was respectively applied. Manual thresholding was only applied when none of the automatized algorithms was able to detect reliably markers without a clear signal to noise ratio, i.e for APP N-terminal, VAMP2, syt1, BACE1 and PS1. Costes method was used as a statistical test to evaluate the correlation between different channels [17]. The p value for all the images tested in this study was equal to 1. The average pixel intensity for each of the proteins was also measured within the surface of the APP accumulation and 15 μm outside by using the ROI manager function and the gray value distribution histogram.

DAB stainings were used to count APP accumulations with an automated macro that was segmenting, counting and measuring the size of each accumulation in different regions of the brain section. Any misdetections were avoided when the user was invited to check for any false-positive counts or by adding any miss-counted accumulations (https://github.com/fabricecordelieres/IJ-Macro_APP-Plaques-NanoZoomer)

Sholl analysis [72] was used to measure the area and number of APP accumulations around amyloid plaques labeled for Aβ42. For this aim, an automated macro was used to isolate and count the number of APP accumulations in intervals of 5 μm, up to 50 μm distance away from plaques in human and mouse sections. The area and number of Aβ42-positive plaques per hippocampal region were also provided by the macro (https://github.com/fabricecordelieres/IJ-Macro_APP-Aggregates-Per-Plaques-NanoZoomer).

In the mouse brain, similar analyses were performed by using the MOC and Sholl analyses. JACop plugin was used to calculate the MOC in individual plaques and APP accumulations. Thresholds were set automatically for each of the markers stained: APP-Cter and APP-Nter (‘Percentil’ algorithm), Aβ1-16 (‘Yen’ algorithm), Methoxy (‘Max entropy’ algorithm). Neuron cell counting was performed at the somatosensory cortex and the molecular layer by using the cell counter plug in of Image J.

### Electron Microscopy - ultrastructure studies

Electron microscopy was performed on samples from the cortex of 2 WT and 3 APP/PS1 mice to visualize the structural content of APP accumulations. To this aim, mice were anesthetized injecting intraperitoneal urethane. After 5 min, mice were perfused intracardially with 35°C oxygenated (saturating carbogen 95% O2 and 5% CO2) ACSF 1X (in mM: 26 NaHCO3, 2.5 KCl, 16.5 glucose, 2.8 pyruvic acid, 0.5 ascorbic acid, 120 NaCl, 1.25 NaH2PO4, 2 CaCl2, 1 MgCl2, pH 7.4, osmolarity of 305 mOsm) for 4 minutes. Afterwards, 2.5 % paraformaldehyde in 0.1 % glutaraldehyde was perfused intracardially for 10 min until animal fixation. Brains were dissected, post-fixed in PFA for 8 hours and sliced coronally at 250 μm. Sections were permeabilized using sodium borohydride 1% (Sigma Aldrich #452882) at RT for 30 min. Sections were then stained with DAB for APP-C and N-terminal or Aβ 1-16.

Brain sections were post-fixed at RT in 1 % osmium tetroxide in the dark. Slices were then dehydrated in an ascending series of ethanol dilutions from 50 to 100 % for 10 min each. Furthermore, slices were incubated into a 50 % mix (vol.) of Epon 812 resin (Electron Microscopy Sciences) and pure ethanol for 2 h at RT and finally in pure resin overnight. After another bath in pure resin for 5 hours, slices were finally embedded between 2 sheets of aclar, and polymerized at 60°C for 48 hours.

The regions of interest were cut at 500 nm thick sections by using an ultramicrotome and a Diatome diamond knife (Leica EM UC7). Sections of 70 nm thick were then collected on copper grids (Electron Microscopy Sciences) and observed with a transmission electron microscope Hitachi H7650 equipped with a Gatan Orius CCD camera.

### Electron Microscopy - Tokuyasu Immunolabeling with gold particles

2 APP/PS1 and 2 WT mice at 9 months were sacrificed and chemically fixed by using the protocol previously described. Sections were then stained with DAB for APP-C or N-terminal. After chemical fixation, small cortical sections were incubated in 2.3M sucrose letting them rinse overnight at RT. Sections were then mounted with pins and immersed in liquid nitrogen to slice them at 70 nm with a 35° angle Diatome diamond knife at −120°C. Sections were collected in copper grids to be washed with water and incubated with PBS 1x – glycine (0.05mM) for 15 min at RT to start the immunolabelling. Sections were first incubated with a blocking solution of goat conjugate for 30 min and then in a PBS1x-0,1% BSA solution in different baths. The primary antibodies used were then added after diluting them in a PBS1x-0,1% BSA solution. The sections were then washed with PBS 1x-0.1% BSA and incubated with the secondary antibody coupled with gold particles (goat anti-rabbit 10 nm with a dilution of 1:40 for APP C-ter and goat anti-mouse 15 nm with a dilution of 1:50 for VAMP2). After six washes of 5 min each with PBS 1x-0,1% BSA, sections were postfixed with 2% glutaraldehyde. The next wash was done in distilled water and a heavy metal contrast incubation was performed in the dark with aqueous uranyl acetate containing 2% methylcellulose for 10 min in ice. Finally, the grids were gently dried using filter paper.

### Statistical analysis

GraphPad Prism 8.0 (GraphPad Software, La Jolla, CA, USA) was used to perform all the statistical analyses. Experimental data were first tested for normal distribution using d’Agostino and Pearson tests. Data were represented using scatter plots in which the mean or median was included or as histograms of the means. Normally distributed data were compared by using a two-tailed t-test and data that were non-normally distributed were compared using a two-tailed Mann-Whitney rank test. Two-way ANOVA was used for analysis with 2 different factors (number of APP accumulations and patient groups) and represented as histograms with error bars (± SEM). Differences between groups were considered statistically significant when p < 0.05.

## Supplementary figures

**Figure S1. Controls and data supporting that full-length APP accumulates in the hippocampus of AD patients** (A, B) Human hippocampal sections stained by DAB against APP-Cter with C1/6.1 antibody ‘A’ or with Y188 ‘B’ and with hematoxylin. The control pictures are realized under the condition of labelling with a preadsorption step of the primary antibody with a recombinant peptide of the APP-Cter corresponding to the epitope. (C) Human hippocampal sections stained by DAB against the APP-Cter with the Y188 antibody. Cases from control, FAD and SAD groups are shown. In controls, APP staining is observed mostly in the soma of neurons in the dentate gyrus granule cell layer (GCL), CA4 and CA3. In addition, staining of the white matter tracts is visible; it is not specific ‘I’. In both FAD and SAD cases, intense extra-somatic APP stainings are observable. (D) Magnifications in the dentate gyrus of pictures as in ‘C’. In the SAD case, intense extra-somatic APP accumulations are observable in the molecular layer (ML), granule cells layer (GCL) and CA4 regions. (E) Histogram of the number of APP-Cter accumulations detected with the C1/6.1 antibody in different hippocampal regions in the control (Ctrl), FAD or SAD conditions. APP accumulations are barely detected in Ctrl whereas they are abundant in AD. (F) Histogram of the area of the APP accumulations detected with C1/6.1 antibody in different hippocampal regions. The CA4 and CA3 regions contain larger APP accumulations. (G) Controls of fluorescent immuno-labeling in human hippocampal sections. In the upper panels, the sections are stained for DAPI, syntaxin1a and Syt1. In the lower panels, only secondary antibodies were incubated in addition to DAPI. The fluorescent immuno-labelings are specific to the primary antibodies. (H, I) Mouse brain sections fluorescently immuno-labeled against APP-Cter with the C1/6.1 antibody ‘H’ or the Y188 antibody ‘I’ and for NeuN and DAPI. In WT, APP is labeled mostly in the soma of the neurons. For example, the different hippocampal layers are detectable in WT. In APP-KO mouse, this neuronal staining is lost. However, with the Y188 antibody, staining of white matter tracts remains in APP-KO revealing its non-specificity in white matter. (J) Human hippocampal sections fluorescently immuno-labeled against APP-Cter with the Y188 and C1/6.1 antibodies. In controls, APP labeled with both Cter antibodies is found mostly in the soma of the neurons. In FAD condition, intense extra-somatic APP stainings, co-labeled with the two antibodies are observable. (K) Close-up picture of staining as in ‘J’ showing APP accumulations co-labelled with the C1/6.1 and Y188 antibodies. (L) Diagram representing the method to calculate the Manders Ovelap Coefficient (MOC) between objects in two different channels. (M) *Left*, Representation of the proportion of overlap between two channels and their calculated MOCs. *Right*, Description of the Costes test. (N) Scatter plot of MOCs between C1/6.1 and Y188 labelings. The analysis reveals a high colocalization between the two channels. (O) *Left*, Representation of an APP accumulation. The intensity of the APP staining is measured within (‘In’) the APP accumulation and normalized to the intensity outside the accumulation (‘Out’), i.e. the surrounding neuropil. *Right*, scatter plot of the intensity of APP-Cter labeling within APP accumulations. The data are normalized to the intensity outside the APP accumulations represented here by a dashed line. The analysis revealed that APP is enriched inside the APP accumulations. (P) Mouse brain sections fluorescently immuno-labeled against APP-Nter with the JRD32 antibody and for NeuN and DAPI. In WT, APP is labelled mostly in the soma of the neurons. In APP-KO mice, this neuronal staining is lost. (Q) Human hippocampal sections fluorescently immuno-labeled against APP-Cter with the Y188 antibody and APP-Nter with the 22C11 antibody. In controls, APP labeled with both Cter and Nter antibodies is mostly found in neuronal soma. In both FAD and SAD cases, intense extra-somatic APP stainings, co-labeled with the two antibodies are observable. (R) Close-up picture of an APP accumulation co-labeled with APP-Cter (Y188) and Nter (22C11). (S) Scatter plot of Manders overlapping coefficients (MOC) (Method described in FigS1L, M) between APP-Cter and Nter labelings. The analysis revealed a high colocalization between the two channels suggesting the presence of full-length APP in the accumulations.

**Figure S2. Controls and data supporting that APP accumulations surround dense-core amyloid plaques** (A, B) Mouse brain sections fluorescently immuno-labeled against Aβ42 with the 12F4 antibody ‘A’ or against Aβ1-16 with the antibody 6E10 ‘B’ and for NeuN and DAPI. In APP/PS1 mice, amyloid plaques are spread all over the brain. They are absent from WT littermate. (C) Staining as in ‘A, B’ but without primary antibody does not display significant fluorescent signal. (D) Human hippocampal sections fluorescently immuno-labelled against APP-Cter with the Y188 antibody and against amyloid plaques with the Aβ1-16 antibody and methoxy-X04. In both FAD and SAD cases, intense Aβ1-16 and methoxy-X04 labelings are observable in the proximity of APP accumulations whereas no staining is observable in control. (E) Close-up pictures of APP accumulations surrounding an amyloid plaque co-labeled with Aβ1-16 and methoxy-X04. The Aβ1-16 and methoxy-X04 stainings colocalize well together but not with the APP-Cter labeling. (F) Scatter plot of MOCs between methoxy-X04 and Aβ1-16 or APP-Cter labelings. The analysis revealed a high colocalization between methoxy-X04 and Aβ-1-16 (amyloid plaques) and a low colocalization between methoxy-X04 and APP-Cter. (G, H) Pictures of a human brain section from an AD case containing the hippocampus and para-hippocampal cortex labeled against APP-Cter, Aβ42 and methoxy-X04. The lower panels in (H) show hippocampal dense-core plaques (revealed by methoxy-X04 and Aβ42) surrounded by APP accumulations. (I, J) Pictures of a human brain section from a control case containing the hippocampus and para-hippocampal cortex labeled against APP-Cter, Aβ1-16 and methoxy-X04. The lower panels in (J) show cortical diffuse plaques labelled by Aβ1-16 but not by methoxy-X04. They are not surrounded by APP accumulations. (K) Pictures as in (I) but labelled for Aβ42. No dense-core amyloid plaques are observable in control cases. (L) Representation of a Sholl analysis to count APP accumulations around an amyloid plaque labeled with Aβ42 in concentric annuli of increasing diameter (intervals of 5 μm), up to 50 μm away from the amyloid plaque. (M) Bland-Altman analysis of the areas covered by APP accumulations and Aβ42-positive amyloid plaques. The difference in areas is plotted in function of the average in areas.

**Figure S3. Controls and data supporting that dense-core plaques are surrounded by BACE1 and contain APP-Nter in their core** (A) Scatter plot of the MOC between methoxy-X04 and BACE1. The analysis reveals a low colocalization between methoxy-X04 and β-secretase. (B) Close-up pictures of hippocampal amyloid plaques labeled at APP-Cter (Y188), Nter (22C11) and methoxy-X04 revealed a remarkable abundance of APP-Nter in the core of dense-core amyloid plaques surrounded by APP accumulations. (C) Scatter plot of the MOC between methoxy-X04 and APP-Nter or Cter. The analysis revealed a high colocalization between methoxy-X04 and APP-Nter (22C11) but not with APP-Cter (Y188) indicating that the colocalization between APP-Nter and the core of the plaques is specific of the Nter domain. (D) Human hippocampal sections immuno-labelled against APP-Cter, PS1 and with methoxy-X04. PS1 is prominently observed in neuronal soma reminiscent of the expression of APP, in addition to a fainter labeling of the neuropil. (E) Close-up pictures of stainings as in ‘D’ showing PS1 and APP accumulations colocalizing in the surrounding of an amyloid plaque labeled with methoxy-X04. (F) Scatter plot of the MOC between APP-Cter and PS1. (G) Scatter plot of the intensity of PS1 labeling within APP accumulations. The data are normalized to the intensity outside the APP accumulations represented here by a dashed line. The analysis revealed that PS1 is modestly enriched inside the APP accumulations. (H) Scatter plot of the MOC between methoxy-X04 and PS1. The analysis reveals a low colocalization between methoxy-X04 and PS1.

**Figure S4. Neuritic markers in hippocampal sections from control cases** (A-E) Human hippocampal sections stained against APP and methoxy-X04 (or DAPI in ‘C’) and with one of the following neuritic marker: MAP2 (A), SMI312 (B), MBP (C), TAU or P-TAU (E). Inlets in the lower left corners display higher magnifications.

**Figure S5. Synaptic vesicle proteins concentrate within APP accumulations in the grey matter** (A, B) Human hippocampal sections fluorescently immuno-labeled against APP-Cter with Y188 and Vamp2 ‘A’ or syntaxin1A (Stx1A) ‘B’. Vamp2 staining is prominent in the neuropil, especially in the axonal terminals of CA4 and CA3; Stx1A staining is bright all over the neuropil. The two presynaptic proteins are present at the level of APP accumulations. (C) Scatter plot of the MOC between methoxy-X04 and synaptic markers (Shank2, Syt1, VAMP2, STX1A). The analysis revealed a low colocalization between methoxy-X04 and these markers. (D) Human hippocampal sections from an AD patient, fluorescently immuno-labeled against APP-Cter (C1/6.1) and myelin basic protein (MBP) enriched in white matter tracts. APP accumulations are mostly observable in MBP-negative regions. (E) Scatter plot of the number of APP accumulations in the white matter (regions MBP-positive) and in the grey matter (regions MBP-negative). The APP accumulations are more abundant in the grey matter. (F) Scatter plot of the area of APP accumulations in the white matter (regions MBP-positive) and in the grey matter (regions MBP-negative). The APP accumulations in the white matter are smaller.

**Figure S6. AD mouse models recapitulate the features of APP accumulations observed in human** (A) Scatter plot of the MOC between APP-Cter and methoxy-X04 or Aβ1-16 orAPP-Nter. The analysis reveals a high colocalization between APP-Cter and Nter but not with Aβ1-16 or methoxy-X04 indicating that amyloid plaques and APP accumulations are non-overlapping. (B) Scatter plot of the MOC between methoxy-X04 and APP-Cter or Aβ1-16 or APP-Nter. The analysis reveals a high colocalization between methoxy-X04 and APP-Nter but not with APP-Cter indicating that amyloid plaques contain APP-Nter but not APP-Cter. (C) Graph of a Sholl analysis of the number of APP accumulations in function of the distance from Aβ1-16 plaques indicating that APP accumulations are more numerous close to the amyloid plaques. (D, F) APP-KI NL/G/F mouse hippocampal sections immuno-labeled against APP-Cter and either Aβ1-16 (D) or APP-Nter (F). (E, G) Close-up pictures of stainings as in ‘D, F’ showing that amyloid plaques containing APP-Nter are surrounded by APP accumulations as in human AD conditions. (H) Close-up pictures of stainings in APP/PS1 showing the presence of VAMP2 in APP accumulations surrounding amyloid plaques labeled with methoxy-X04.

## Bibliography

1. Arai H, Lee VM, Otvos L, Greenberg BD, Lowery DE, Sharma SK, Schmidt ML, Trojanowski JQ (1990) Defined neurofilament, tau, and beta-amyloid precursor protein epitopes distinguish Alzheimer from non-Alzheimer senile plaques. PNAS 87:2249–2253

2. Barthet G, Georgakopoulos A, Robakis NK (2012) Cellular mechanisms of γ-secretase substrate selection, processing and toxicity. Prog Neurobiol 98:166–175. doi: 10.1016/j.pneurobio.2012.05.006

3. Barthet G, Jordà-Siquier T, Rumi-Masante J, Bernadou F, Müller U, Mulle C (2018) Presenilin-mediated cleavage of APP regulates synaptotagmin-7 and presynaptic plasticity. Nat Commun 9:4780. doi: 10.1038/s41467-018-06813-x

4. Barthet G, Mulle C (2020) Presynaptic failure in Alzheimer’s disease. Prog Neurobiol 101801. doi: 10.1016/j.pneurobio.2020.101801

5. Barthet G, Shioi J, Shao Z, Ren Y, Georgakopoulos A, Robakis NK (2011) Inhibitors of γ-secretase stabilize the complex and differentially affect processing of amyloid precursor protein and other substrates. FASEB J 25:2937–2946. doi: 10.1096/fj.11-183806

6. Baumkötter F, Schmidt N, Vargas C, Schilling S, Weber R, Wagner K, Fiedler S, Klug W, Radzimanowski J, Nickolaus S, Keller S, Eggert S, Wild K, Kins S (2014) Amyloid precursor protein dimerization and synaptogenic function depend on copper binding to the growth factor-like domain. J Neurosci 34:11159–11172. doi: 10.1523/JNEUROSCI.0180-14.2014

7. Bentahir M, Nyabi O, Verhamme J, Tolia A, Horré K, Wiltfang J, Esselmann H, De Strooper B (2006) Presenilin clinical mutations can affect gamma-secretase activity by different mechanisms. J Neurochem 96:732–742. doi: 10.1111/j.1471-4159.2005.03578.x

8. Bolte S, Cordelières FP (2006) A guided tour into subcellular colocalization analysis in light microscopy. J Microsc 224:213–232. doi: 10.1111/j.1365-2818.2006.01706.x

9. Cacquevel M, Aeschbach L, Houacine J, Fraering PC (2012) Alzheimer’s disease-linked mutations in presenilin-1 result in a drastic loss of activity in purified γ-secretase complexes. PLoS ONE 7:e35133. doi: 10.1371/journal.pone.0035133

10. Cras P, Kawai M, Lowery D, Gonzalez-DeWhitt P, Greenberg B, Perry G (1991) Senile plaque neurites in Alzheimer disease accumulate amyloid precursor protein. Proc Natl Acad Sci USA 88:7552–7556. doi: 10.1073/pnas.88.17.7552

11. Cummings BJ, Su JH, Geddes JW, Van Nostrand WE, Wagner SL, Cunningham DD, Cotman CW (1992) Aggregation of the amyloid precursor protein within degenerating neurons and dystrophic neurites in Alzheimer’s disease. Neuroscience 48:763–777. doi: 10.1016/0306-4522(92)90265-4

12. D’Andrea MR, Nagele RG (2002) MAP-2 immunolabeling can distinguish diffuse from dense-core amyloid plaques in brains with Alzheimer’s disease. Biotech Histochem 77:95–103

13. De Strooper B, Saftig P, Craessaerts K, Vanderstichele H, Guhde G, Annaert W, Von Figura K, Van Leuven F (1998) Deficiency of presenilin-1 inhibits the normal cleavage of amyloid precursor protein. Nature 391:387–390. doi: 10.1038/34910

14. DeKosky ST, Scheff SW (1990) Synapse loss in frontal cortex biopsies in Alzheimer’s disease: correlation with cognitive severity. Ann Neurol 27:457–464. doi: 10.1002/ana.410270502

15. Delaère P, He Y, Fayet G, Duyckaerts C, Hauw JJ (1993) Beta A4 deposits are constant in the brain of the oldest old: an immunocytochemical study of 20 French centenarians. Neurobiol Aging 14:191–194. doi: 10.1016/0197-4580(93)90096-t

16. Deyts C, Clutter M, Herrera S, Jovanovic N, Goddi A, Parent AT (2016) Loss of presenilin function is associated with a selective gain of APP function. Elife 5. doi: 10.7554/eLife.15645

17. Dunn KW, Kamocka MM, McDonald JH (2011) A practical guide to evaluating colocalization in biological microscopy. Am J Physiol, Cell Physiol 300:C723–742. doi: 10.1152/ajpcell.00462.2010

18. Eisele YS, Obermüller U, Heilbronner G, Baumann F, Kaeser SA, Wolburg H, Walker LC, Staufenbiel M, Heikenwalder M, Jucker M (2010) Peripherally applied Abeta-containing inoculates induce cerebral beta-amyloidosis. Science 330:980–982. doi: 10.1126/science.1194516

19. Ferguson B, Matyszak MK, Esiri MM, Perry VH (1997) Axonal damage in acute multiple sclerosis lesions. Brain 120 (Pt 3):393–399. doi: 10.1093/brain/120.3.393

20. Gentleman SM, Nash MJ, Sweeting CJ, Graham DI, Roberts GW (1993) Beta-amyloid precursor protein (beta APP) as a marker for axonal injury after head injury. Neurosci Lett 160:139–144. doi: 10.1016/0304-3940(93)90398-5

21. Georgakopoulos A, Marambaud P, Efthimiopoulos S, Shioi J, Cui W, Li HC, Schütte M, Gordon R, Holstein GR, Martinelli G, Mehta P, Friedrich VL, Robakis NK (1999) Presenilin-1 forms complexes with the cadherin/catenin cell-cell adhesion system and is recruited to intercellular and synaptic contacts. Mol Cell 4:893–902. doi: 10.1016/s1097-2765(00)80219-1

22. Glenner GG, Wong CW (1984) Alzheimer’s disease and Down’s syndrome: sharing of a unique cerebrovascular amyloid fibril protein. Biochem Biophys Res Commun 122:1131–1135. doi: 10.1016/0006-291x(84)91209-9

23. Goate A, Chartier-Harlin MC, Mullan M, Brown J, Crawford F, Fidani L, Giuffra L, Haynes A, Irving N, James L (1991) Segregation of a missense mutation in the amyloid precursor protein gene with familial Alzheimer’s disease. Nature 349:704–706. doi: 10.1038/349704a0

24. Gouras GK, Almeida CG, Takahashi RH (2005) Intraneuronal Abeta accumulation and origin of plaques in Alzheimer’s disease. Neurobiol Aging 26:1235–1244. doi: 10.1016/j.neurobiolaging.2005.05.022

25. Gunawardena S, Yang G, Goldstein LSB (2013) Presenilin controls kinesin-1 and dynein function during APP-vesicle transport in vivo. Hum Mol Genet 22:3828–3843. doi: 10.1093/hmg/ddt237

26. Hadley KC, Rakhit R, Guo H, Sun Y, Jonkman JE, McLaurin J, Hazrati L-N, Emili A, Chakrabartty A (2015) Determining composition of micron-scale protein deposits in neurodegenerative disease by spatially targeted optical microproteomics. eLife 4. doi: 10.7554/eLife.09579

27. Hardy JA, Higgins GA (1992) Alzheimer’s disease: the amyloid cascade hypothesis. Science 256:184–185. doi: 10.1126/science.1566067

28. Haytural H, Mermelekas G, Emre C, Nigam SM, Carroll SL, Winblad B, Bogdanovic N, Barthet G, Granholm A-C, Orre LM, Tjernberg LO, Frykman S (2020) The Proteome of the Dentate Terminal Zone of the Perforant Path Indicates Presynaptic Impairment in Alzheimer Disease. Mol Cell Proteomics 19:128–141. doi: 10.1074/mcp.RA119.001737

29. Honer WG, Barr AM, Sawada K, Thornton AE, Morris MC, Leurgans SE, Schneider JA, Bennett DA (2012) Cognitive reserve, presynaptic proteins and dementia in the elderly. Transl Psychiatry 2:e114. doi: 10.1038/tp.2012.38

30. Huang L-K, Chao S-P, Hu C-J (2020) Clinical trials of new drugs for Alzheimer disease. Journal of Biomedical Science 27:18. doi: 10.1186/s12929-019-0609-7

31. Huang Y-WA, Zhou B, Wernig M, Südhof TC (2017) ApoE2, ApoE3, and ApoE4 Differentially Stimulate APP Transcription and Aβ Secretion. Cell 168:427–441.e21. doi: 10.1016/j.cell.2016.12.044

32. Jankowsky JL, Fadale DJ, Anderson J, Xu GM, Gonzales V, Jenkins NA, Copeland NG, Lee MK, Younkin LH, Wagner SL, Younkin SG, Borchelt DR (2004) Mutant presenilins specifically elevate the levels of the 42 residue beta-amyloid peptide in vivo: evidence for augmentation of a 42-specific gamma secretase. Hum Mol Genet 13:159–170. doi: 10.1093/hmg/ddh019

33. Jiang Y, Mullaney KA, Peterhoff CM, Che S, Schmidt SD, Boyer-Boiteau A, Ginsberg SD, Cataldo AM, Mathews PM, Nixon RA (2010) Alzheimer’s-related endosome dysfunction in Down syndrome is Abeta-independent but requires APP and is reversed by BACE-1 inhibition. Proc Natl Acad Sci USA 107:1630–1635. doi: 10.1073/pnas.0908953107

34. Kandalepas PC, Sadleir KR, Eimer WA, Zhao J, Nicholson DA, Vassar R (2013) The Alzheimer’s β-secretase BACE1 localizes to normal presynaptic terminals and to dystrophic presynaptic terminals surrounding amyloid plaques. Acta Neuropathol 126:329–352. doi: 10.1007/s00401-013-1152-3

35. Kapur JN, Sahoo PK, Wong AKC (1985) A new method for gray-level picture thresholding using the entropy of the histogram. Computer Vision, Graphics, and Image Processing 29:273–285. doi: 10.1016/0734-189X(85)90125-2

36. Katzman R, Terry R, DeTeresa R, Brown T, Davies P, Fuld P, Renbing X, Peck A (1988) Clinical, pathological, and neurochemical changes in dementia: a subgroup with preserved mental status and numerous neocortical plaques. Ann Neurol 23:138–144. doi: 10.1002/ana.410230206

37. Kohli BM, Pflieger D, Mueller LN, Carbonetti G, Aebersold R, Nitsch RM, Konietzko U (2012) Interactome of the amyloid precursor protein APP in brain reveals a protein network involved in synaptic vesicle turnover and a close association with Synaptotagmin-1. J Proteome Res 11:4075–4090. doi: 10.1021/pr300123g

38. Koo EH, Sisodia SS, Archer DR, Martin LJ, Weidemann A, Beyreuther K, Fischer P, Masters CL, Price DL (1990) Precursor of amyloid protein in Alzheimer disease undergoes fast anterograde axonal transport. Proc Natl Acad Sci USA 87:1561–1565

39. Kuhlmann T, Lingfeld G, Bitsch A, Schuchardt J, Brück W (2002) Acute axonal damage in multiple sclerosis is most extensive in early disease stages and decreases over time. Brain 125:2202–2212. doi: 10.1093/brain/awf235

40. Kwart D, Gregg A, Scheckel C, Murphy EA, Paquet D, Duffield M, Fak J, Olsen O, Darnell RB, Tessier-Lavigne M (2019) A Large Panel of Isogenic APP and PSEN1 Mutant Human iPSC Neurons Reveals Shared Endosomal Abnormalities Mediated by APP β-CTFs, Not Aβ. Neuron 104:256–270.e5. doi: 10.1016/j.neuron.2019.07.010

41. Laßek M, Weingarten J, Einsfelder U, Brendel P, Müller U, Volknandt W (2013) Amyloid precursor proteins are constituents of the presynaptic active zone. J Neurochem 127:48–56. doi: 10.1111/jnc.12358

42. Lazarov O, Morfini GA, Pigino G, Gadadhar A, Chen X, Robinson J, Ho H, Brady ST, Sisodia SS (2007) Impairments in fast axonal transport and motor neuron deficits in transgenic mice expressing familial Alzheimer’s disease-linked mutant presenilin 1. J Neurosci 27:7011–7020. doi: 10.1523/JNEUROSCI.4272-06.2007

43. Lee J-H, Yu WH, Kumar A, Lee S, Mohan PS, Peterhoff CM, Wolfe DM, Martinez-Vicente M, Massey AC, Sovak G, Uchiyama Y, Westaway D, Cuervo AM, Nixon RA (2010) Lysosomal proteolysis and autophagy require presenilin 1 and are disrupted by Alzheimer-related PS1 mutations. Cell 141:1146–1158. doi: 10.1016/j.cell.2010.05.008

44. Lemere CA, Blusztajn JK, Yamaguchi H, Wisniewski T, Saido TC, Selkoe DJ (1996) Sequence of deposition of heterogeneous amyloid beta-peptides and APO E in Down syndrome: implications for initial events in amyloid plaque formation. Neurobiol Dis 3:16–32. doi: 10.1006/nbdi.1996.0003

45. Lundgren JL, Ahmed S, Schedin-Weiss S, Gouras GK, Winblad B, Tjernberg LO, Frykman S (2015) ADAM10 and BACE1 are localized to synaptic vesicles. J Neurochem 135:606–615. doi: 10.1111/jnc.13287

46. Mangialasche F, Solomon A, Winblad B, Mecocci P, Kivipelto M (2010) Alzheimer’s disease: clinical trials and drug development. Lancet Neurol 9:702–716. doi: 10.1016/S1474-4422(10)70119-8

47. Mann DM (1988) Alzheimer’s disease and Down’s syndrome. Histopathology 13:125–137. doi: 10.1111/j.1365-2559.1988.tb02018.x

48. Martinez–Vicente M, Talloczy Z, Wong E, Tang G, Koga H, Kaushik S, de Vries R, Arias E, Harris S, Sulzer D, Cuervo A (2010) CARGO RECOGNITION FAILURE IS RESPONSIBLE FOR INEFFICIENT AUTOPHAGY IN HUNTINGTON’S DISEASE. Nat Neurosci 13:567–576. doi: 10.1038/nn.2528

49. Masliah E, Mallory M, Hansen L, Alford M, DeTeresa R, Terry R (1993) An antibody against phosphorylated neurofilaments identifies a subset of damaged association axons in Alzheimer’s disease. Am J Pathol 142:871–882

50. Mayer MC, Schauenburg L, Thompson-Steckel G, Dunsing V, Kaden D, Voigt P, Schaefer M, Chiantia S, Kennedy TE, Multhaup G (2016) Amyloid precursor-like protein 1 (APLP1) exhibits stronger zinc-dependent neuronal adhesion than amyloid precursor protein and APLP2. J Neurochem 137:266–276. doi: 10.1111/jnc.13540

51. McInnes J, Wierda K, Snellinx A, Bounti L, Wang Y-C, Stancu I-C, Apóstolo N, Gevaert K, Dewachter I, Spires-Jones TL, De Strooper B, De Wit J, Zhou L, Verstreken P (2018) Synaptogyrin-3 Mediates Presynaptic Dysfunction Induced by Tau. Neuron 97:823–835.e8. doi: 10.1016/j.neuron.2018.01.022

52. Mesulam MM, Geula C (1994) Butyrylcholinesterase reactivity differentiates the amyloid plaques of aging from those of dementia. Ann Neurol 36:722–727. doi: 10.1002/ana.410360506

53. Meyer-Luehmann M, Coomaraswamy J, Bolmont T, Kaeser S, Schaefer C, Kilger E, Neuenschwander A, Abramowski D, Frey P, Jaton AL, Vigouret J-M, Paganetti P, Walsh DM, Mathews PM, Ghiso J, Staufenbiel M, Walker LC, Jucker M (2006) Exogenous induction of cerebral beta-amyloidogenesis is governed by agent and host. Science 313:1781–1784. doi: 10.1126/science.1131864

54. Mori I, Goshima F, Mizuno T, Imai Y, Kohsaka S, Ito H, Koide N, Yoshida T, Yokochi T, Kimura Y, Nishiyama Y (2005) Axonal injury in experimental herpes simplex encephalitis. Brain Res 1057:186–190. doi: 10.1016/j.brainres.2005.07.037

55. Motter R, Vigo-Pelfrey C, Kholodenko D, Barbour R, Johnson-Wood K, Galasko D, Chang L, Miller B, Clark C, Green R (1995) Reduction of beta-amyloid peptide42 in the cerebrospinal fluid of patients with Alzheimer’s disease. Ann Neurol 38:643–648. doi: 10.1002/ana.410380413

56. Munter L-M, Voigt P, Harmeier A, Kaden D, Gottschalk KE, Weise C, Pipkorn R, Schaefer M, Langosch D, Multhaup G (2007) GxxxG motifs within the amyloid precursor protein transmembrane sequence are critical for the etiology of Abeta42. EMBO J 26:1702–1712. doi: 10.1038/sj.emboj.7601616

57. Narang HK (1980) High-resolution electron microscopic analysis of the amyloid fibril in Alzheimer’s disease. Journal of Neuropathology and Experimental Neurology 39:621–631. doi: 10.1097/00005072-198011000-00001

58. Neely KM, Green KN, LaFerla FM (2011) Presenilin is necessary for efficient proteolysis through the autophagy-lysosome system in a γ-secretase-independent manner. J Neurosci 31:2781–2791. doi: 10.1523/JNEUROSCI.5156-10.2010

59. Nixon RA, Wegiel J, Kumar A, Yu WH, Peterhoff C, Cataldo A, Cuervo AM (2005) Extensive involvement of autophagy in Alzheimer disease: an immuno-electron microscopy study. J Neuropathol Exp Neurol 64:113–122. doi: 10.1093/jnen/64.2.113

60. Peters F, Salihoglu H, Pratsch K, Herzog E, Pigoni M, Sgobio C, Lichtenthaler SF, Neumann U, Herms J (2019) Tau deletion reduces plaque-associated BACE1 accumulation and decelerates plaque formation in a mouse model of Alzheimer’s disease. EMBO J 38:e102345. doi: 10.15252/embj.2019102345

61. Rorke-Adams LB (2011) 45 - Neuropathology of Abusive Head Trauma. In: Jenny C (ed) Child Abuse and Neglect. W.B. Saunders, Philadelphia, pp 413–428

62. Ross CA, Poirier MA (2004) Protein aggregation and neurodegenerative disease. Nature Medicine 10:S10–S17. doi: 10.1038/nm1066

63. Rovelet-Lecrux A, Hannequin D, Raux G, Le Meur N, Laquerrière A, Vital A, Dumanchin C, Feuillette S, Brice A, Vercelletto M, Dubas F, Frebourg T, Campion D (2006) APP locus duplication causes autosomal dominant early-onset Alzheimer disease with cerebral amyloid angiopathy. Nat Genet 38:24–26. doi: 10.1038/ng1718

64. Sadleir KR, Kandalepas PC, Buggia-Prévot V, Nicholson DA, Thinakaran G, Vassar R (2016) Presynaptic dystrophic neurites surrounding amyloid plaques are sites of microtubule disruption, BACE1 elevation, and increased Aβ generation in Alzheimer’s disease. Acta Neuropathol 132:235–256. doi: 10.1007/s00401-016-1558-9

65. Saito T, Matsuba Y, Mihira N, Takano J, Nilsson P, Itohara S, Iwata N, Saido TC (2014) Single App knock-in mouse models of Alzheimer’s disease. Nat Neurosci 17:661–663. doi: 10.1038/nn.3697

66. Scheff SW, Price DA, Schmitt FA, DeKosky ST, Mufson EJ (2007) Synaptic alterations in CA1 in mild Alzheimer disease and mild cognitive impairment. Neurology 68:1501–1508. doi: 10.1212/01.wnl.0000260698.46517.8f

67. Selkoe DJ (2002) Alzheimer’s disease is a synaptic failure. Science 298:789–791. doi: 10.1126/science.1074069

68. Shahmoradian SH, Lewis AJ, Genoud C, Hench J, Moors TE, Navarro PP, Castaño-Díez D, Schweighauser G, Graff-Meyer A, Goldie KN, Sütterlin R, Huisman E, Ingrassia A, Gier Y de, Rozemuller AJM, Wang J, Paepe AD, Erny J, Staempfli A, Hoernschemeyer J, Großerüschkamp F, Niedieker D, El-Mashtoly SF, Quadri M, Van IJcken WFJ, Bonifati V, Gerwert K, Bohrmann B, Frank S, Britschgi M, Stahlberg H, Van de Berg WDJ, Lauer ME (2019) Lewy pathology in Parkinson’s disease consists of crowded organelles and lipid membranes. Nat Neurosci 22:1099–1109. doi: 10.1038/s41593-019-0423-2

69. Sharoar MG, Hu X, Ma X-M, Zhu X, Yan R (2019) Sequential formation of different layers of dystrophic neurites in Alzheimer’s brains. Molecular Psychiatry 24:1369–1382. doi: 10.1038/s41380-019-0396-2

70. Sherrington R, Rogaev EI, Liang Y, Rogaeva EA, Levesque G, Ikeda M, Chi H, Lin C, Li G, Holman K, Tsuda T, Mar L, Foncin JF, Bruni AC, Montesi MP, Sorbi S, Rainero I, Pinessi L, Nee L, Chumakov I, Pollen D, Brookes A, Sanseau P, Polinsky RJ, Wasco W, Da Silva HA, Haines JL, Perkicak-Vance MA, Tanzi RE, Roses AD, Fraser PE, Rommens JM, St George-Hyslop PH (1995) Cloning of a gene bearing missense mutations in early-onset familial Alzheimer’s disease. Nature 375:754–760. doi: 10.1038/375754a0

71. Shoji M, Hirai S, Yamaguchi H, Harigaya Y, Kawarabayashi T (1990) Amyloid beta-protein precursor accumulates in dystrophic neurites of senile plaques in Alzheimer-type dementia. Brain Res 512:164–168. doi: 10.1016/0006-8993(90)91187-l

72. Sholl DA (1953) Dendritic organization in the neurons of the visual and motor cortices of the cat. J Anat 87:387–406

73. Sisodia SS, Koo EH, Hoffman PN, Perry G, Price DL (1993) Identification and transport of full-length amyloid precursor proteins in rat peripheral nervous system. J Neurosci 13:3136–3142

74. Sleegers K, Brouwers N, Gijselinck I, Theuns J, Goossens D, Wauters J, Del-Favero J, Cruts M, van Duijn CM, Van Broeckhoven C (2006) APP duplication is sufficient to cause early onset Alzheimer’s dementia with cerebral amyloid angiopathy. Brain 129:2977–2983. doi: 10.1093/brain/awl203

75. Soba P, Eggert S, Wagner K, Zentgraf H, Siehl K, Kreger S, Löwer A, Langer A, Merdes G, Paro R, Masters CL, Müller U, Kins S, Beyreuther K (2005) Homo- and heterodimerization of APP family members promotes intercellular adhesion. EMBO J 24:3624–3634. doi: 10.1038/sj.emboj.7600824

76. Soto C, Pritzkow S (2018) Protein misfolding, aggregation, and conformational strains in neurodegenerative diseases. Nature Neuroscience 21:1332–1340. doi: 10.1038/s41593-018-0235-9

77. Su JH, Cummings BJ, Cotman CW (1994) Early phosphorylation of tau in Alzheimer’s disease occurs at Ser-202 and is preferentially located within neurites. Neuroreport 5:2358–2362. doi: 10.1097/00001756-199411000-00037

78. Sun L, Zhou R, Yang G, Shi Y (2017) Analysis of 138 pathogenic mutations in presenilin-1 on the in vitro production of Aβ42 and Aβ40 peptides by γ-secretase. Proc Natl Acad Sci USA 114:E476–E485. doi: 10.1073/pnas.1618657114

79. Terry RD, Masliah E, Salmon DP, Butters N, DeTeresa R, Hill R, Hansen LA, Katzman R (1991) Physical basis of cognitive alterations in Alzheimer’s disease: synapse loss is the major correlate of cognitive impairment. Ann Neurol 30:572–580. doi: 10.1002/ana.410300410

80. Umehara F, Abe M, Koreeda Y, Izumo S, Osame M (2000) Axonal damage revealed by accumulation of beta-amyloid precursor protein in HTLV-I-associated myelopathy. J Neurol Sci 176:95–101. doi: 10.1016/s0022-510x(00)00324-5

81. Weingarten MD, Lockwood AH, Hwo SY, Kirschner MW (1975) A protein factor essential for microtubule assembly. Proc Natl Acad Sci USA 72:1858–1862. doi: 10.1073/pnas.72.5.1858

82. de Wilde MC, Overk CR, Sijben JW, Masliah E (2016) Meta-analysis of synaptic pathology in Alzheimer’s disease reveals selective molecular vesicular machinery vulnerability. Alzheimers Dement 12:633–644. doi: 10.1016/j.jalz.2015.12.005

83. Wilhelm BG, Mandad S, Truckenbrodt S, Kröhnert K, Schäfer C, Rammner B, Koo SJ, Claßen GA, Krauss M, Haucke V, Urlaub H, Rizzoli SO (2014) Composition of isolated synaptic boutons reveals the amounts of vesicle trafficking proteins. Science 344:1023–1028. doi: 10.1126/science.1252884

84. Wisniewski HM, Wegiel J, Wang KC, Kujawa M, Lach B (1989) Ultrastructural studies of the cells forming amyloid fibers in classical plaques. The Canadian Journal of Neurological Sciences Le Journal Canadien Des Sciences Neurologiques 16:535–542. doi: 10.1017/s0317167100029887

85. Woodruff G, Young JE, Martinez FJ, Buen F, Gore A, Kinaga J, Li Z, Yuan SH, Zhang K, Goldstein LSB (2013) The presenilin-1 ΔE9 mutation results in reduced γ-secretase activity, but not total loss of PS1 function, in isogenic human stem cells. Cell Rep 5:974–985. doi: 10.1016/j.celrep.2013.10.018

86. Yen JC, Chang FJ, Chang S (1995) A new criterion for automatic multilevel thresholding. IEEE Trans Image Process 4:370–378. doi: 10.1109/83.366472

87. Yu L, Tasaki S, Schneider JA, Arfanakis K, Duong DM, Wingo AP, Wingo TS, Kearns N, Thatcher GRJ, Seyfried NT, Levey AI, De Jager PL, Bennett DA (2020) Cortical Proteins Associated With Cognitive Resilience in Community-Dwelling Older Persons. JAMA Psychiatry. doi: 10.1001/jamapsychiatry.2020.1807

88. Zhang X, Garbett K, Veeraraghavalu K, Wilburn B, Gilmore R, Mirnics K, Sisodia SS (2012) A role for presenilins in autophagy revisited: normal acidification of lysosomes in cells lacking PSEN1 and PSEN2. J Neurosci 32:8633–8648. doi: 10.1523/JNEUROSCI.0556-12.2012

89. Zhang X-M, Cai Y, Xiong K, Cai H, Luo X-G, Feng J-C, Clough RW, Struble RG, Patrylo PR, Yan X-X (2009) Beta-secretase-1 elevation in transgenic mouse models of Alzheimer’s disease is associated with synaptic/axonal pathology and amyloidogenesis: implications for neuritic plaque development. Eur J Neurosci 30:2271–2283. doi: 10.1111/j.1460-9568.2009.07017.x

90. Zhou L, McInnes J, Wierda K, Holt M, Herrmann AG, Jackson RJ, Wang Y-C, Swerts J, Beyens J, Miskiewicz K, Vilain S, Dewachter I, Moechars D, De Strooper B, Spires-Jones TL, De Wit J, Verstreken P (2017) Tau association with synaptic vesicles causes presynaptic dysfunction. Nat Commun 8:15295. doi: 10.1038/ncomms15295

