## Supplementary material for "APP accumulates around dense-core amyloid plaques with presynaptic proteins in Alzheimer’s disease brain": Table 1

Table 1 - Clinical resume table

| Case | Gender | Braak stage | Age of onset | Age of death | PMD (h) | Brain weight (g) | CSF PH | Brain bank | Number of patients |
|---|---|---|---|---|---|---|---|---|---|
| Control | F/M (73/27%) | I-III | - | 79,6 | 8,54 | 1204 | 6,75 | NBB/KI | 11 |
| SAD | F/M (92/8%) | V-VI | 72,1 | 81 | 11,08 | 1061 | 6,43 | NBB/KI | 12 |
| FAD | F/M (50/50%) | V-VI | 40,7 | 40,7 | 3,5 | 1101 | 6,22 | NBB/KI | 4 |

M – male, F – female, KI – Karolinska Institutet, NBB – Netherlands Brain Bank
