## Supplementary Information for "APP accumulates around dense-core amyloid plaques with presynaptic proteins in Alzheimer’s disease brain"

Table S1-3

Table S1- Clinical data from control patients

| Control case number | Gender | Braak stage | Age of onset | Age of death | APOE | PMD (h) | Brain weight (g) | CSF PH | Brain bank |
|---|---|---|---|---|---|---|---|---|---|
| 1 | F | I | - | 73 | 4:4 | 7 | 1444 | - | NBB |
| 2 | F | II | - | 76 | 3:3 | 7 | 1072 | 6,87 | NBB |
| 3 | F | III | - | 81 | 3:3 | 5 | 1192 | 6,77 | NBB |
| 4 | F | II | - | 84 | 3:3 | 5 | 1027 | 6,68 | NBB |
| 5 | F | II | - | 83 | 4:4 | 6 | 1030 | 6,6 | NBB |
| 6 | F | II | - | 82 | 3:3 | 3 | 1221 | 6,34 | NBB |
| 7 | F | II | - | 84 | 3:3 | 6 | 1266 | 7,65 | NBB |
| 8 | F | II | - | 93 | 3:3 | 5 | 1145 | - | NBB |
| 9 | M | - | - | 80 | 3:4 | 16 | 1200 | - | KI |
| 10 | M | - | - | 69 | 3:3 | 22 | 1350 | - | KI |
| 11 | M | - | - | 71 | 3:3 | 12 | 1300 | 6,40 | KI |
| Average | F/M (73/27%) | - | | 79,6 | - | 8,54 | 1204 | 6,75 | |

M–male, F– female, KI–Karolinska Institutet, NBB–Netherlands Brain Bank

Table S2- Clinical data from SAD patients

| SAD case number | Gender | Braak stage | Age of onset | Age of death | APOE | PMD (h) | Brain weight (g) | CSF PH | Brain bank |
|---|---|---|---|---|---|---|---|---|---|
| 1 | F | V | 79 | 85 | 4:4 | 5 | 919 | 6,35 | NBB |
| 2 | F | VI | 87 | 90 | 3:3 | 4 | 1045 | 6,39 | NBB |
| 3 | F | V | 63 | 70 | 3:3 | 5 | 1245 | 6,73 | NBB |
| 4 | F | V | 73 | 91 | 4:4 | 6 | 1196 | 6,01 | NBB |
| 5 | F | VI | 56 | 70 | 3:3 | 6 | 824 | 6 | NBB |
| 6 | F | VI | 79 | 80 | 4:4 | 4 | 1112 | 6,26 | NBB |
| 7 | F | VI | 53 | 70 | 4:4 | 4 | 894 | 6,42 | NBB |
| 8 | F | VI | 78 | 86 | 3:3 | 5 | 1266 | 7,65 | NBB |
| 9 | F | V | 77 | 82 | 4:4 | 4 | 999 | 6,08 | NBB |
| 10 | F | VI | - | 78 | 3:4 | 12 | 1080 | - | KI |
| 11 | M | V | 65 | 71 | 3:4 | 12 | 1235 | - | KI |
| 12 | F | V | 84 | 99 | - | 66 | 923 | - | KI |
| Average | F/M ( (92/8%) | - | 72,1 | 81 | - | 11,08 | 1061 | 6,43 | |

M–male, F–female, KI–Karolinska Institutet, NBB–Netherlands Brain Bank

Table S3 - Clinical data from FAD patients

| FAD case number | Gender | Braak stage | Age of onset | Age of death | APOE | PMD (h) | Brain weight (g) | CSF PH | Brain bank |
|---|---|---|---|---|---|---|---|---|---|
| PS1-G206A | F | VI | 35 | 43 | 3:3 | 2 | 845 | 6,22 | NBB |
| PS1-I143T | M | VI | 37 | 45 | 3:3 | - | 1225 | - | KI |
| PS1-I143T | F | VI | 36 | 43 | 3:3 | - | - | - | KI |
| PS1-H163Y | M | V | 55 | 63 | 2:4 | 5 | 1234 | - | KI |
| Average | F/M (50/50%) | - | 40,7 | 48,5 | - | 3,5 | 1101 | 6,22 | |

M–male, F–female, KI–Karolinska Institutet, NBB–Netherlands Brain Bank

### Supplementary figures

Figure S1

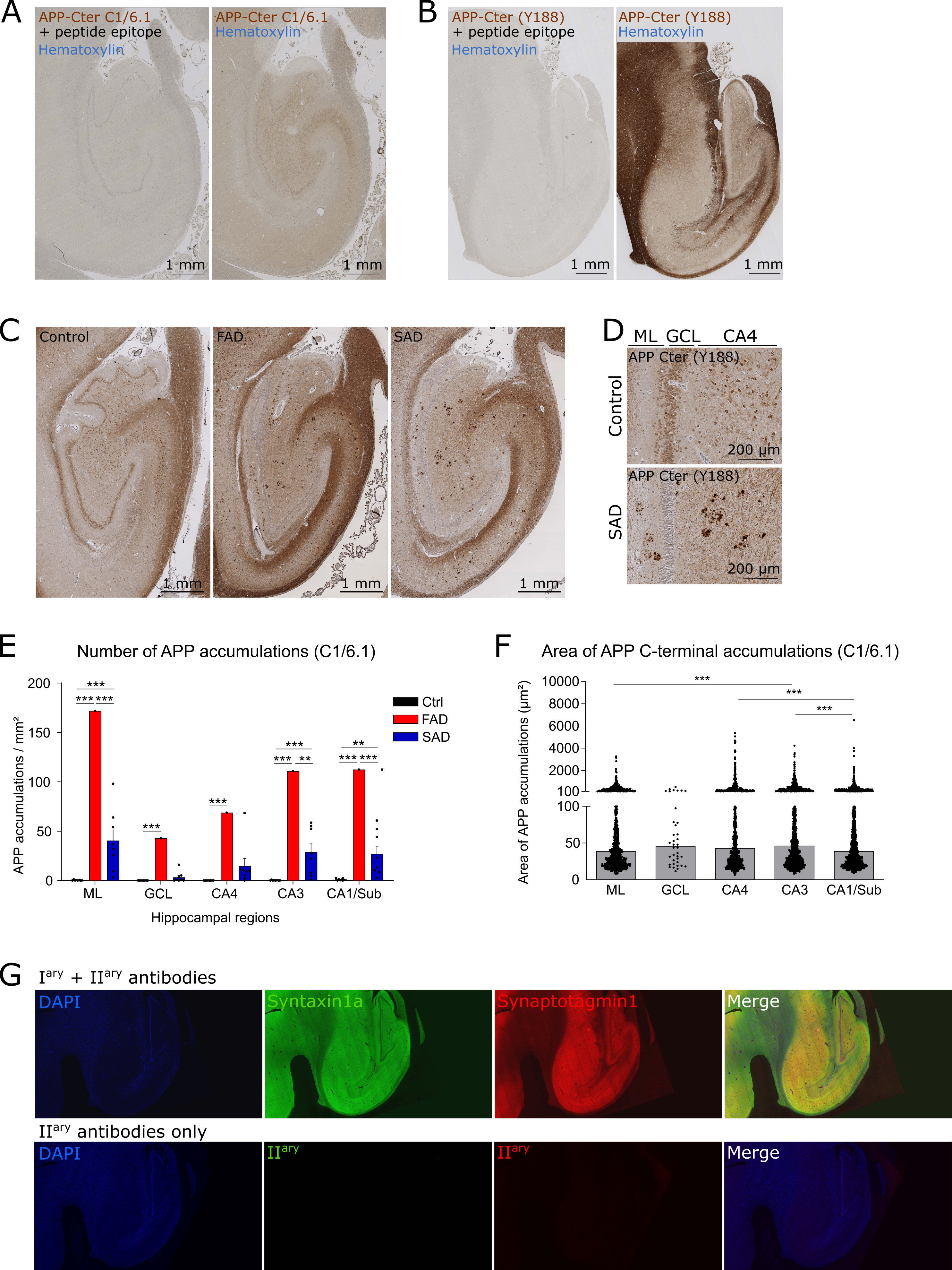

A
APP-Cter C1/6.1
+ peptide epitope
Hematoxylin
APP-Cter C1/6.1
Hematoxylin
1 mm
B
APP-Cter (Y188)
+ peptide epitope
Hematoxylin
APP-Cter (Y188)
Hematoxylin
1 mm
C
Control
FAD
SAD
1 mm
D
ML GCL CA4
Control
SAD
APP Cter (Y188)
200 µm
E
Number of APP accumulations (C1/6.1)
APP accumulations / mm²
Ctrl
FAD
SAD
ML GCL CA4 CA3 CA1/Sub
Hippocampal regions
F
Area of APP C-terminal accumulations (C1/6.1)
Area of APP accumulations (µm²)
ML GCL CA4 CA3 CA1/Sub
G
I^ary + II^ary antibodies
DAPI
Syntaxin1a
Synaptotagmin1
Merge
II^ary antibodies only
II^ary

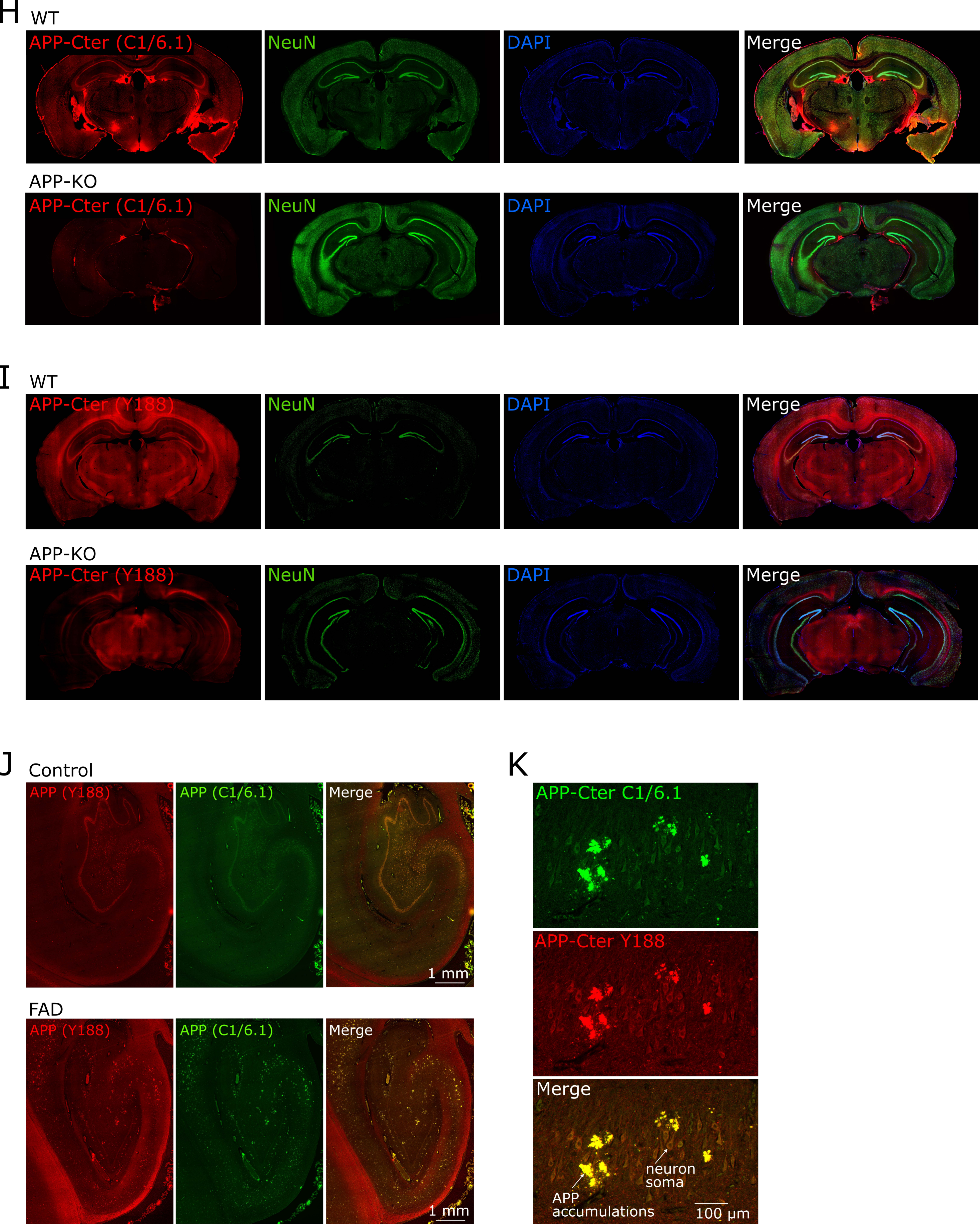

H
WT
APP-Cter (C1/6.1)
NeuN
DAPI
Merge
APP-KO
APP-Cter (C1/6.1)
NeuN
DAPI
Merge
I
WT
APP-Cter (Y188)
NeuN
DAPI
Merge
APP-KO
APP-Cter (Y188)
NeuN
DAPI
Merge
J
Control
APP (Y188)
APP (C1/6.1)
Merge
1 mm
FAD
APP (Y188)
APP (C1/6.1)
Merge
1 mm
K
APP-Cter C1/6.1
APP-Cter Y188
Merge
neuron soma
APP accumulations
100 µm

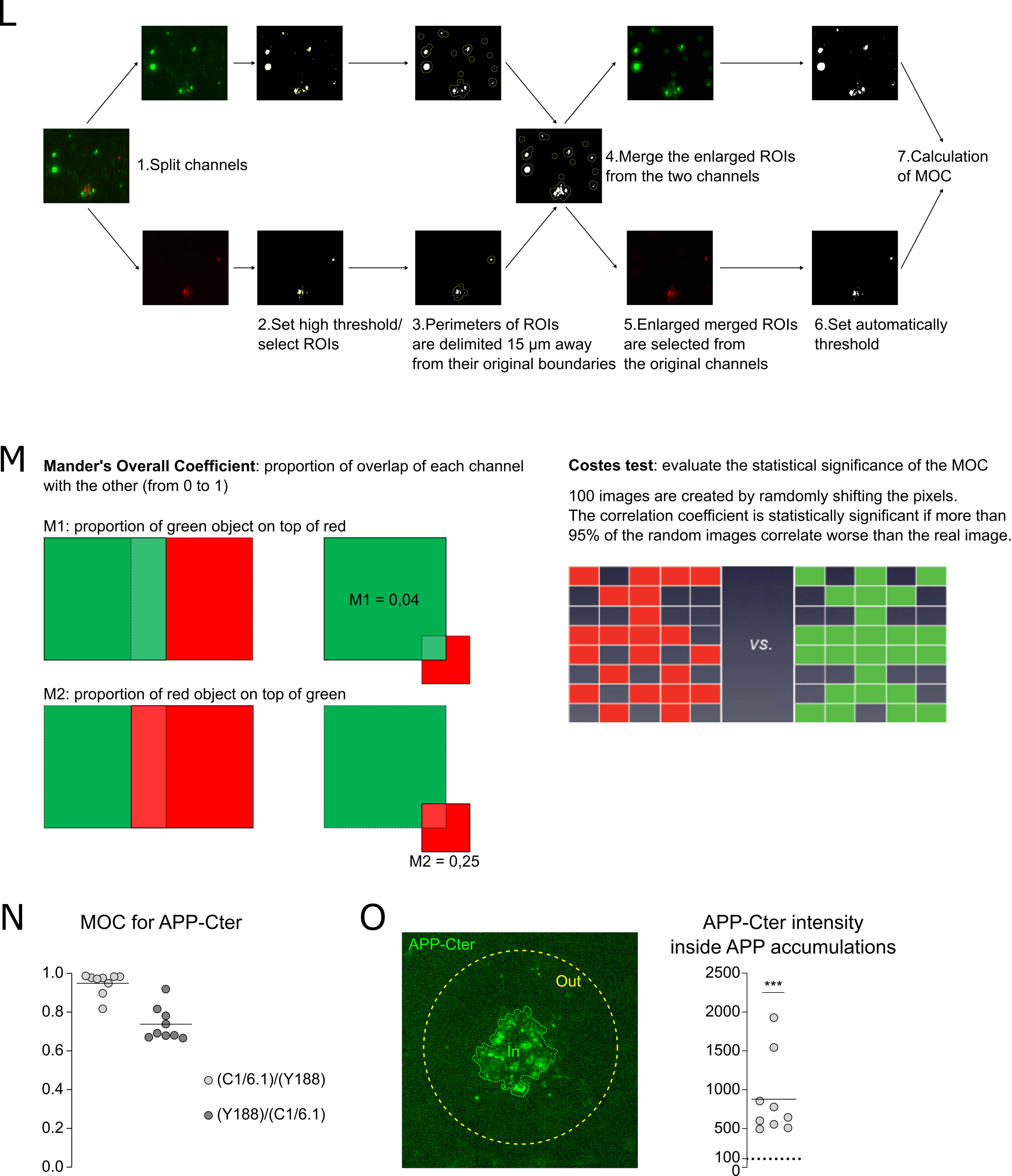
L
1.Split channels
2.Set high threshold/ select ROIs
3.Perimeters of ROIs are delimited 15 µm away from their original boundaries
4.Merge the enlarged ROIs from the two channels
5.Enlarged merged ROIs are selected from the original channels
6.Set automatically threshold
7.Calculation of MOC
M
Mander's Overall Coefficient: proportion of overlap of each channel with the other (from 0 to 1)
M1: proportion of green object on top of red
M1 = 0,04
M2: proportion of red object on top of green
M2 = 0,25
Costes test: evaluate the statistical significance of the MOC
100 images are created by ramdomly shifting the pixels.
The correlation coefficient is statistically significant if more than 95% of the random images correlate worse than the real image.
vs.
N
MOC for APP-Cter
1.0
0.8
0.6
0.4
0.2
0.0
(C1/6.1)/(Y188)
(Y188)/(C1/6.1)
O
APP-Cter
Out
In
APP-Cter intensity inside APP accumulations
2500
2000
1500
1000
500
100
0
***

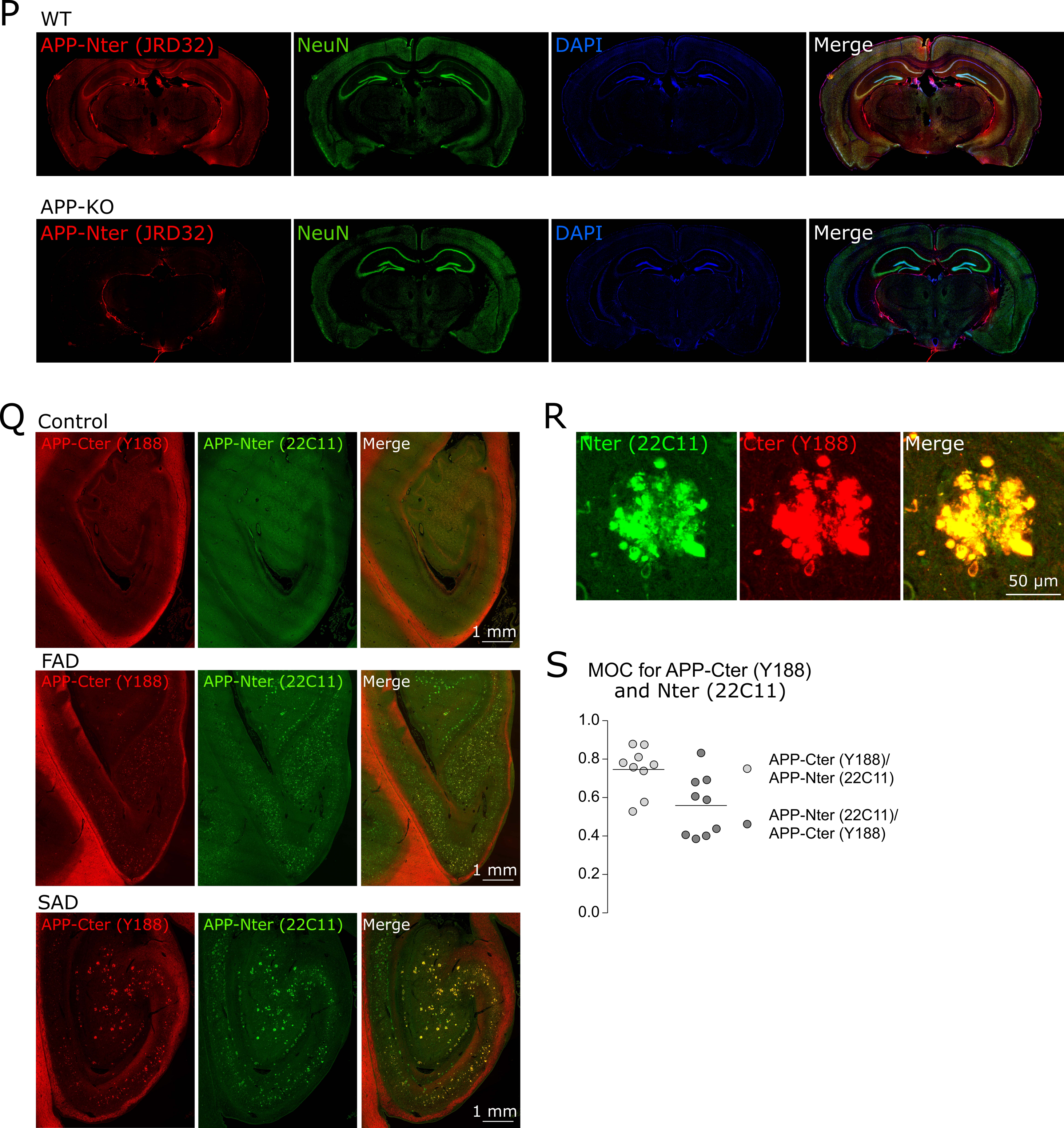
P
WT
APP-Nter (JRD32)
NeuN
DAPI
Merge
APP-KO
APP-Nter (JRD32)
NeuN
DAPI
Merge
Q
Control
APP-Cter (Y188)
APP-Nter (22C11)
Merge
1 mm
FAD
APP-Cter (Y188)
APP-Nter (22C11)
Merge
1 mm
SAD
APP-Cter (Y188)
APP-Nter (22C11)
Merge
1 mm
R
Nter (22C11)
Cter (Y188)
Merge
50 μm
S
MOC for APP-Cter (Y188)
and Nter (22C11)
1.0
0.8
0.6
0.4
0.2
0.0
APP-Cter (Y188)/
APP-Nter (22C11)
APP-Nter (22C11)/
APP-Cter (Y188)

Figure S2

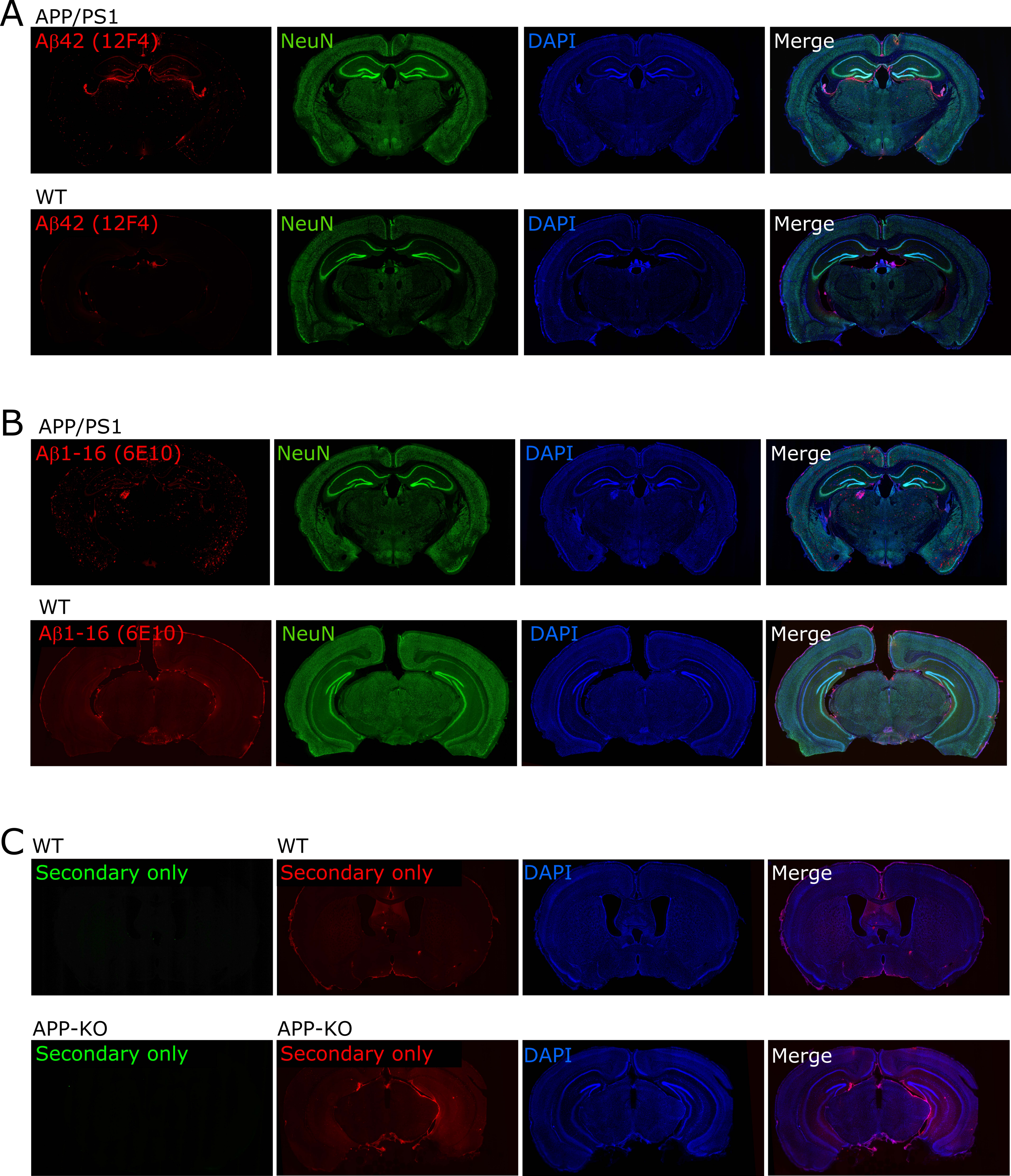
A
APP/PS1
Aβ42 (12F4)
NeuN
DAPI
Merge
WT
Aβ42 (12F4)
NeuN
DAPI
Merge
B
APP/PS1
Aβ1-16 (6E10)
NeuN
DAPI
Merge
WT
Aβ1-16 (6E10)
NeuN
DAPI
Merge
C
WT
Secondary only
WT
Secondary only
DAPI
Merge
APP-KO
Secondary only
APP-KO
Secondary only
DAPI
Merge

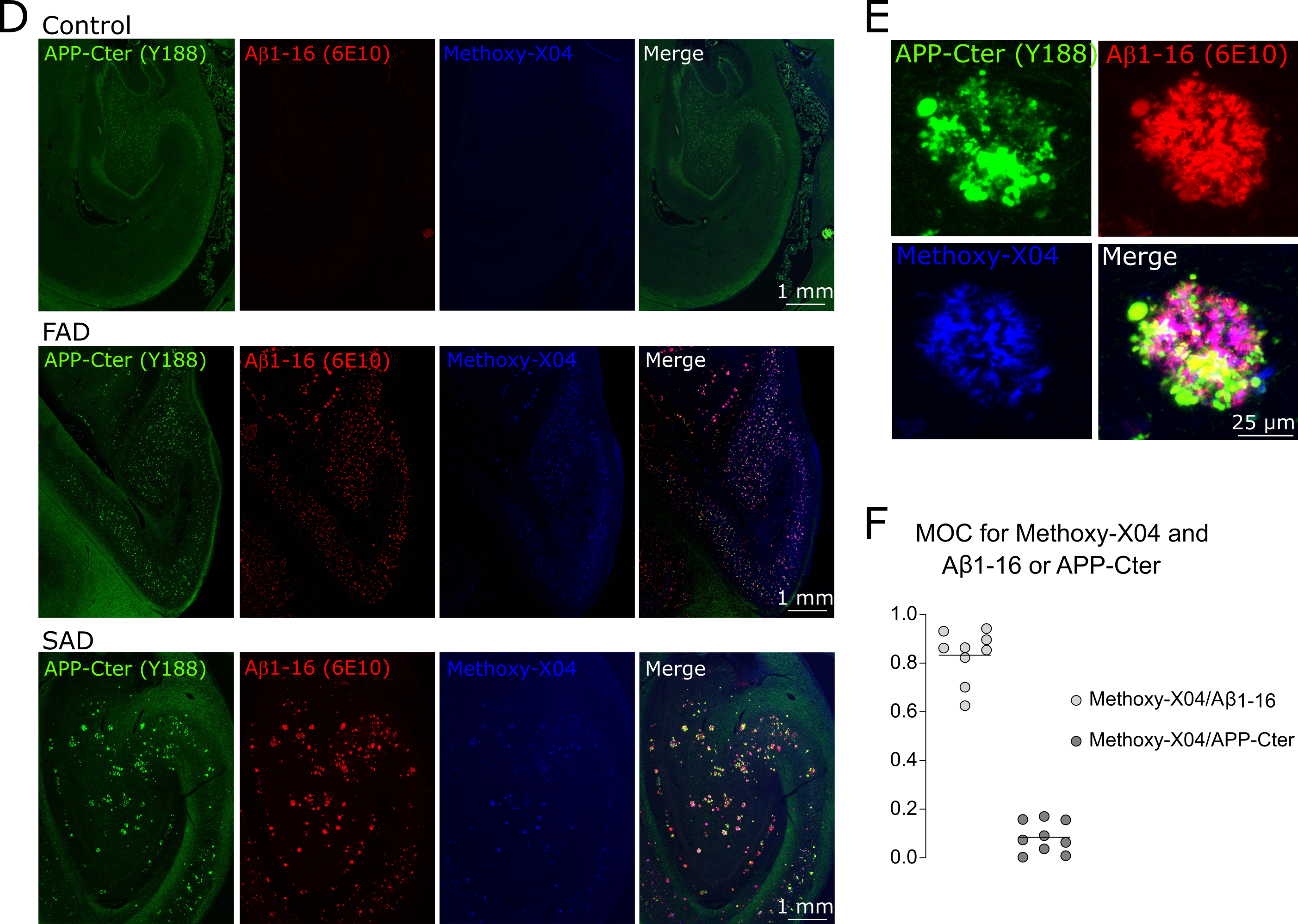
D
Control
APP-Cter (Y188)
Aβ1-16 (6E10)
Methoxy-X04
Merge
1 mm
FAD
APP-Cter (Y188)
Aβ1-16 (6E10)
Methoxy-X04
Merge
1 mm
SAD
APP-Cter (Y188)
Aβ1-16 (6E10)
Methoxy-X04
Merge
1 mm
E
APP-Cter (Y188)
Aβ1-16 (6E10)
Methoxy-X04
Merge
25 μm
F
MOC for Methoxy-X04 and
Aβ1-16 or APP-Cter
1.0
0.8
0.6
0.4
0.2
0.0
Methoxy-X04/Aβ1-16
Methoxy-X04/APP-Cter

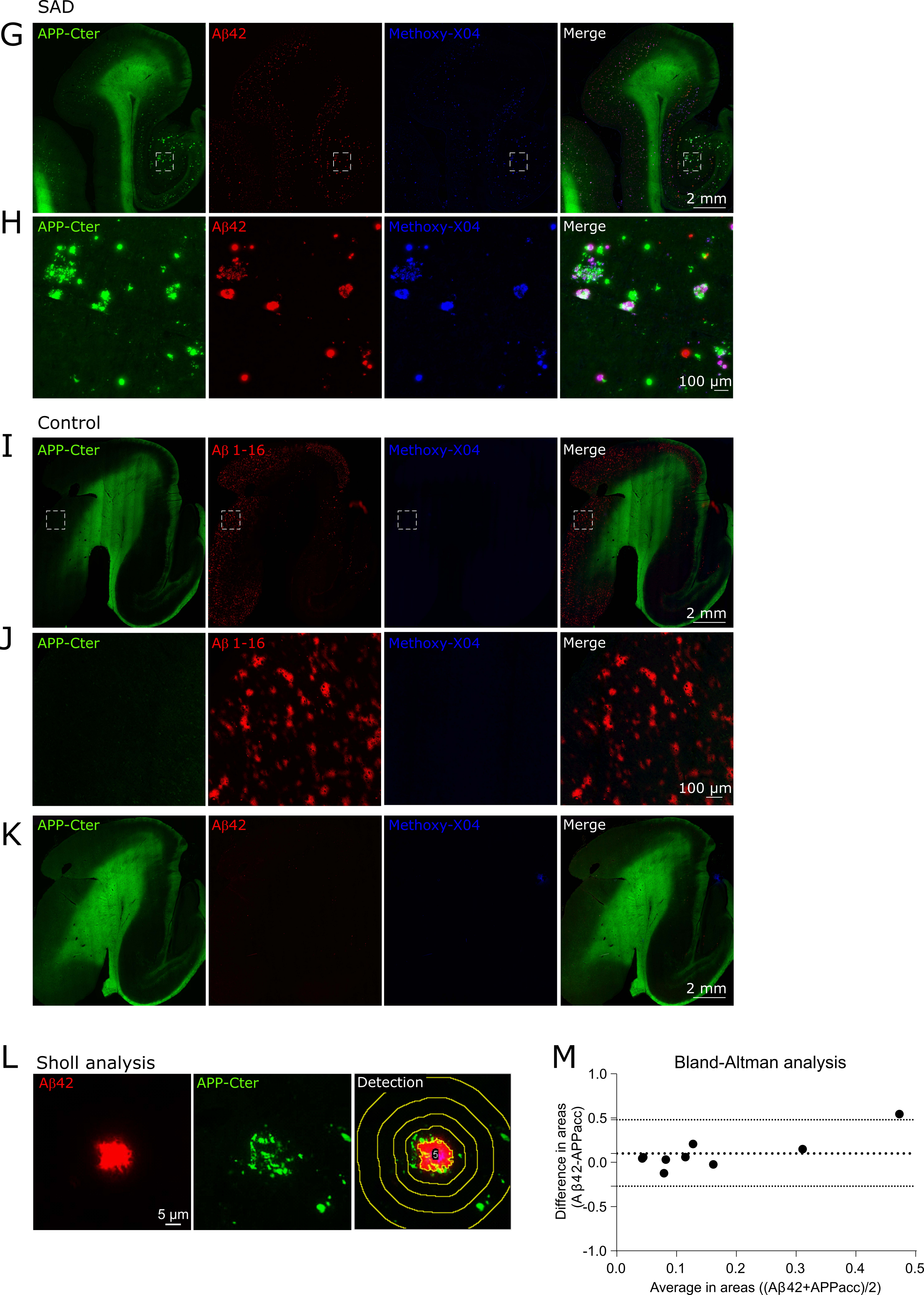

SAD
G
APP-Cter
Aβ42
Methoxy-X04
Merge
2 mm
H
APP-Cter
Aβ42
Methoxy-X04
Merge
100 µm
Control
I
APP-Cter
Aβ 1-16
Methoxy-X04
Merge
2 mm
J
APP-Cter
Aβ 1-16
Methoxy-X04
Merge
100 µm
K
APP-Cter
Aβ42
Methoxy-X04
Merge
2 mm
L
Sholl analysis
Aβ42
APP-Cter
Detection
5 µm
M
Bland-Altman analysis
Difference in areas (Aβ42-APPacc)
1.0
0.5
0.0
-0.5
-1.0
Average in areas ((Aβ42+APPacc)/2)
0.0
0.1
0.2
0.3
0.4
0.5

Figure S3

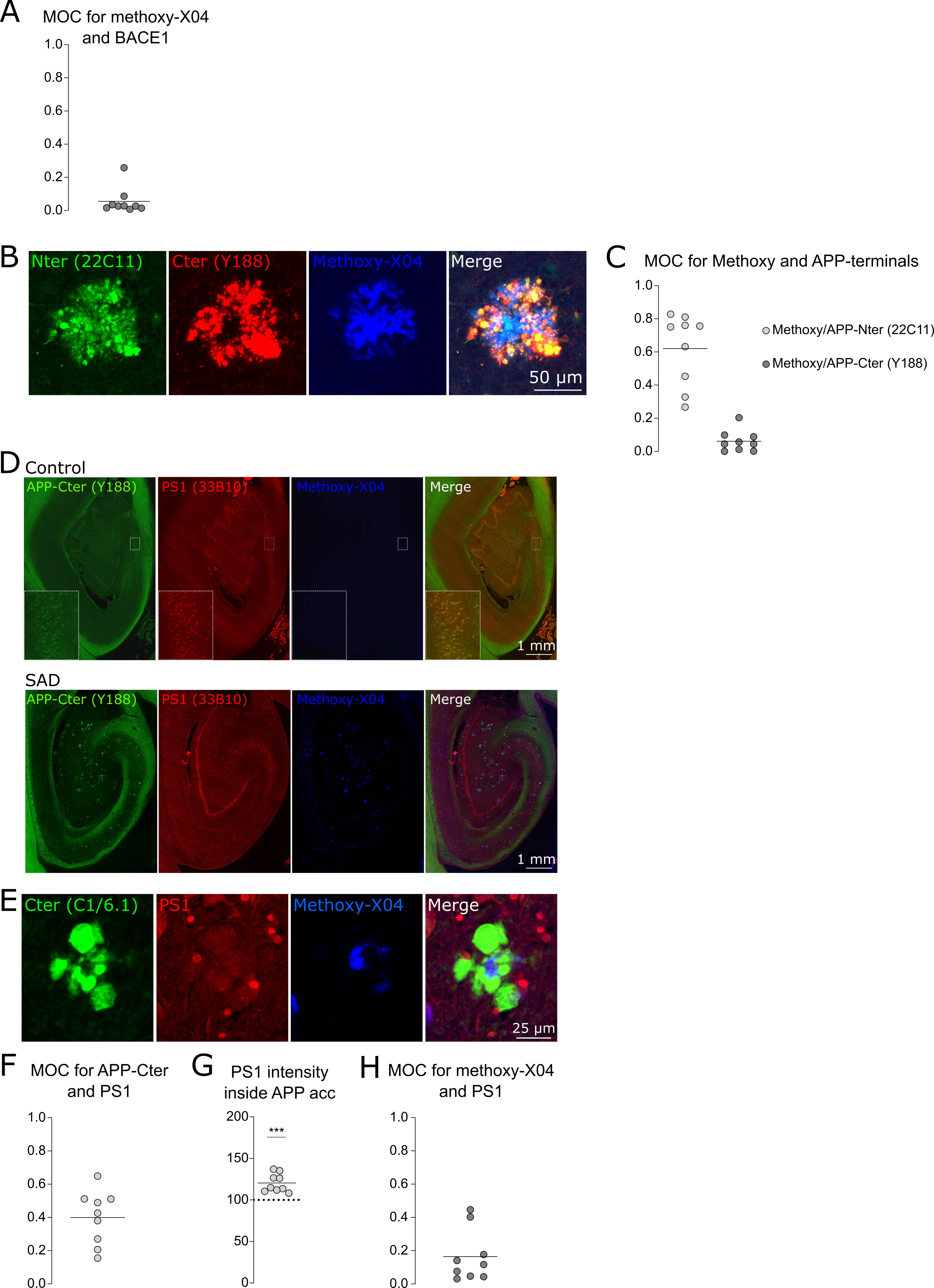

A
MOC for methoxy-X04 and BACE1
1.0
0.8
0.6
0.4
0.2
0.0
B
Nter (22C11)
Cter (Y188)
Methoxy-X04
Merge
50 µm
C
MOC for Methoxy and APP-terminals
Methoxy/APP-Nter (22C11)
Methoxy/APP-Cter (Y188)
D
Control
APP-Cter (Y188)
PS1 (33B10)
Methoxy-X04
Merge
1 mm
SAD
E
Cter (C1/6.1)
PS1
25 µm
F
MOC for APP-Cter and PS1
G
PS1 intensity inside APP acc
200
150
100
50
0
***
H
MOC for methoxy-X04 and PS1

Figure S4

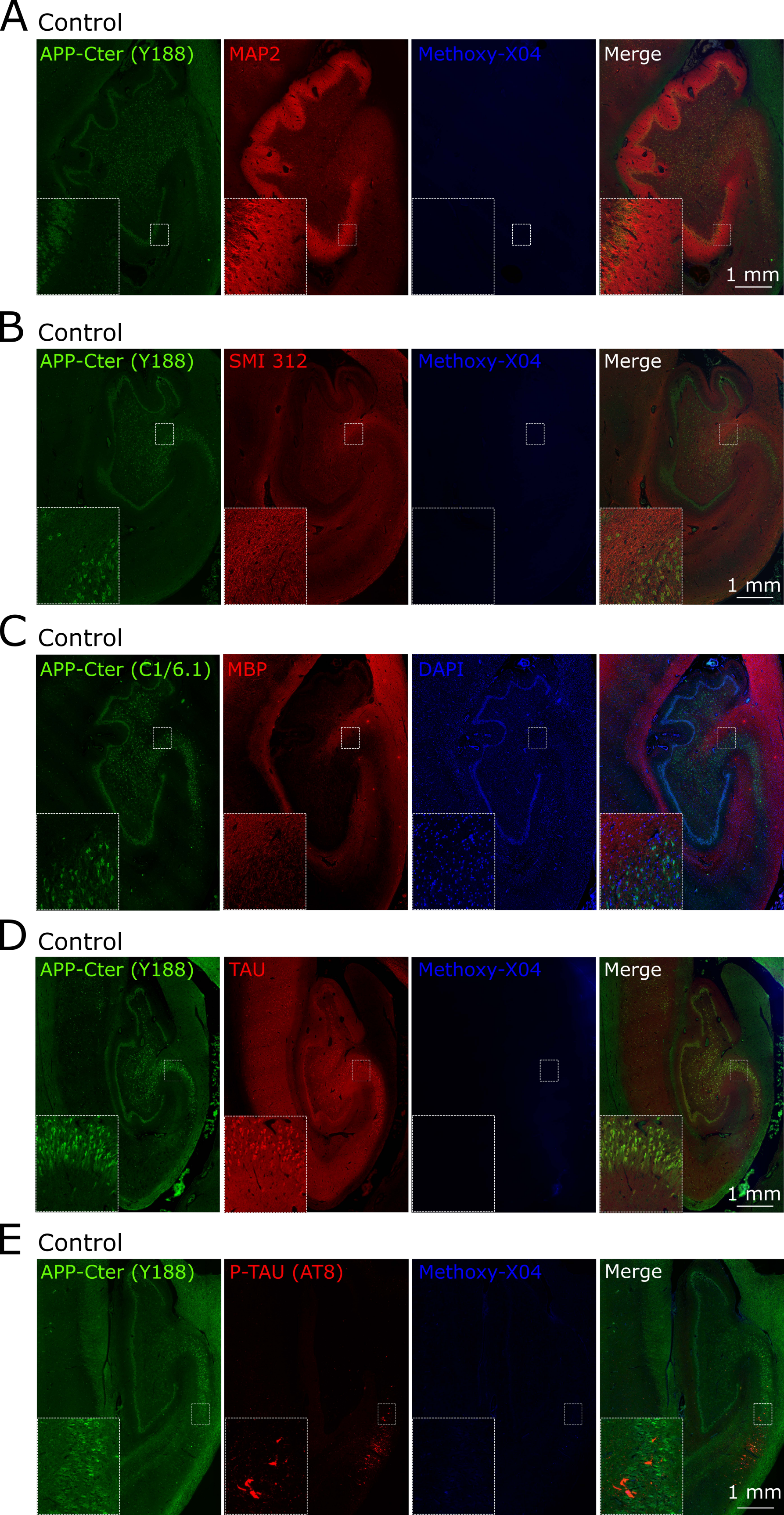
A
Control
APP-Cter (Y188)
MAP2
Methoxy-X04
Merge
1 mm
B
Control
APP-Cter (Y188)
SMI 312
Methoxy-X04
Merge
1 mm
C
Control
APP-Cter (C1/6.1)
MBP
DAPI
D
Control
APP-Cter (Y188)
TAU
Methoxy-X04
Merge
1 mm
E
Control
APP-Cter (Y188)
P-TAU (AT8)
Methoxy-X04
Merge
1 mm

Figure S5

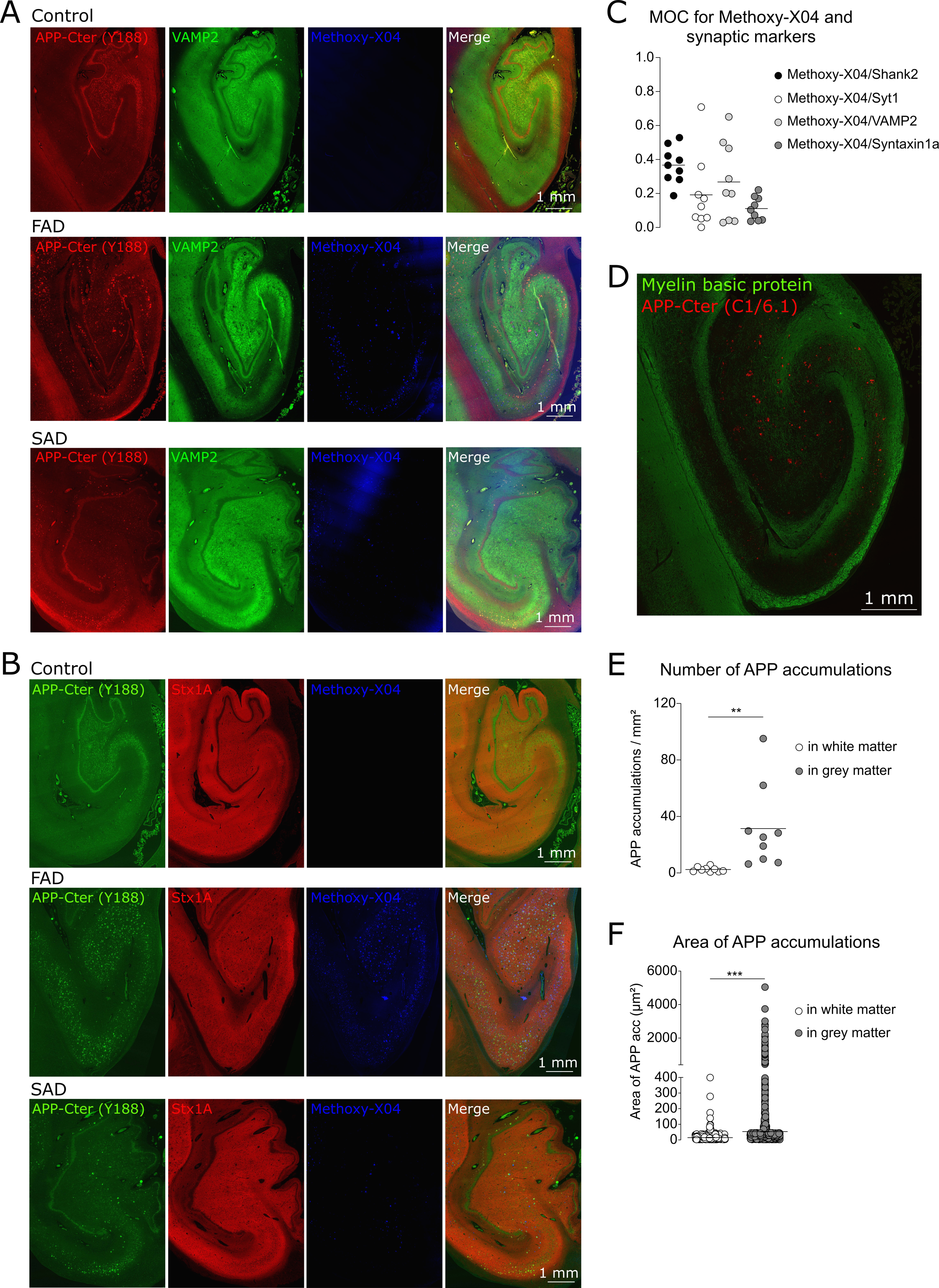

A Control
APP-Cter (Y188)
VAMP2
Methoxy-X04
Merge
1 mm
FAD
SAD
B Control
APP-Cter (Y188)
Stx1A
Methoxy-X04
Merge
FAD
SAD
C MOC for Methoxy-X04 and synaptic markers
1.0
0.8
0.6
0.4
0.2
0.0
Methoxy-X04/Shank2
Methoxy-X04/Syt1
Methoxy-X04/VAMP2
Methoxy-X04/Syntaxin1a
D Myelin basic protein
APP-Cter (C1/6.1)
1 mm
E Number of APP accumulations
APP accumulations / mm²
120
80
40
0
**
in white matter
in grey matter
F Area of APP accumulations
Area of APP acc (µm²)
6000
4000
2000
400
300
200
100
0
***
in white matter
in grey matter

Figure S6

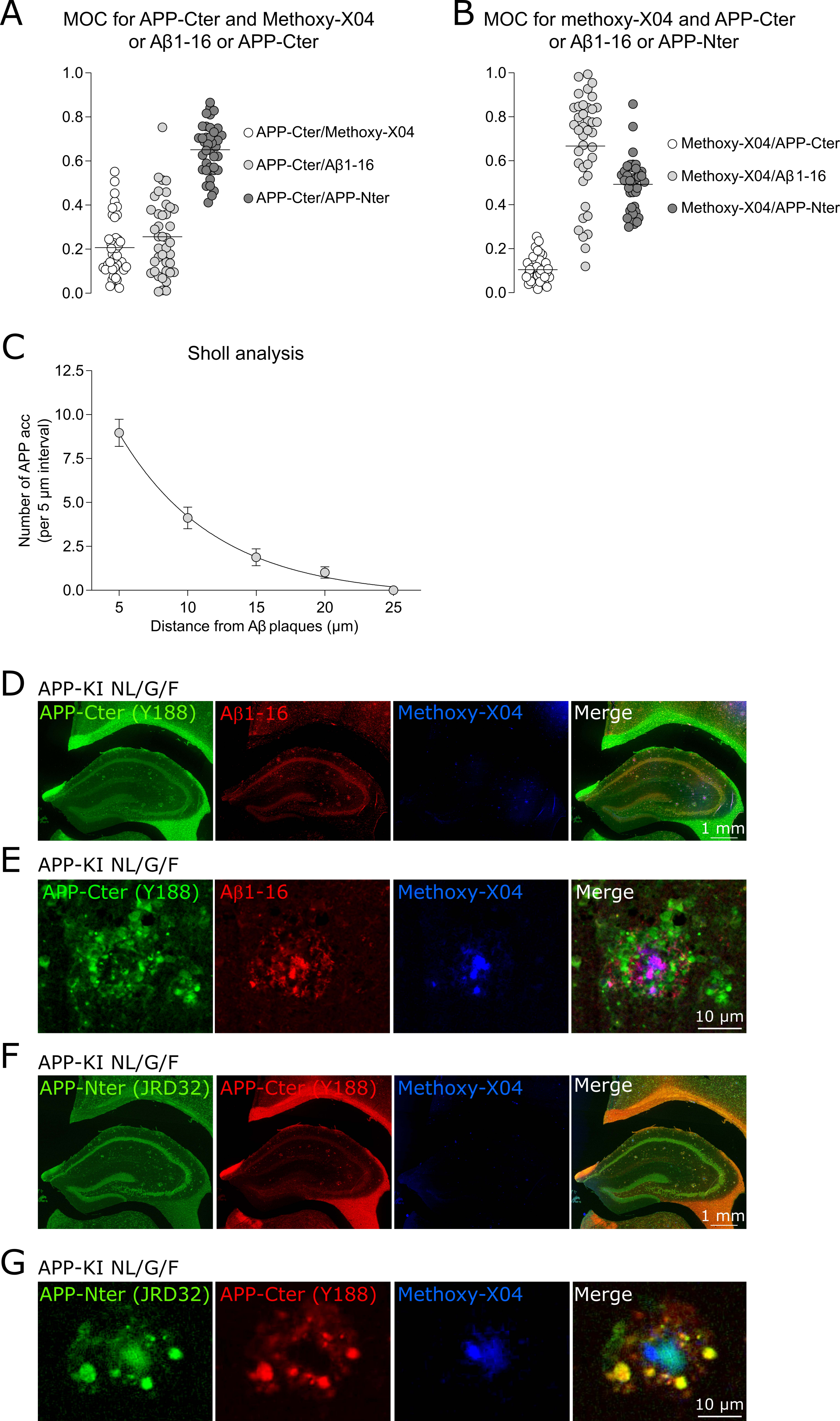

A
MOC for APP-Cter and Methoxy-X04
or Aβ1-16 or APP-Cter
1.0
0.8
0.6
0.4
0.2
0.0
APP-Cter/Methoxy-X04
APP-Cter/Aβ1-16
APP-Cter/APP-Nter
B
MOC for methoxy-X04 and APP-Cter
or Aβ1-16 or APP-Nter
1.0
0.8
0.6
0.4
0.2
0.0
Methoxy-X04/APP-Cter
Methoxy-X04/Aβ1-16
Methoxy-X04/APP-Nter
C
Sholl analysis
12.5
10.0
7.5
5.0
2.5
0.0
Number of APP acc
(per 5 µm interval)
5
10
15
20
25
Distance from Aβ plaques (µm)
D
APP-KI NL/G/F
APP-Cter (Y188)
Aβ1-16
Methoxy-X04
Merge
1 mm
E
APP-KI NL/G/F
APP-Cter (Y188)
Aβ1-16
Methoxy-X04
Merge
10 µm
F
APP-KI NL/G/F
APP-Nter (JRD32)
APP-Cter (Y188)
Methoxy-X04
Merge
1 mm
G
APP-KI NL/G/F
APP-Nter (JRD32)
APP-Cter (Y188)
Methoxy-X04
Merge
10 µm

H

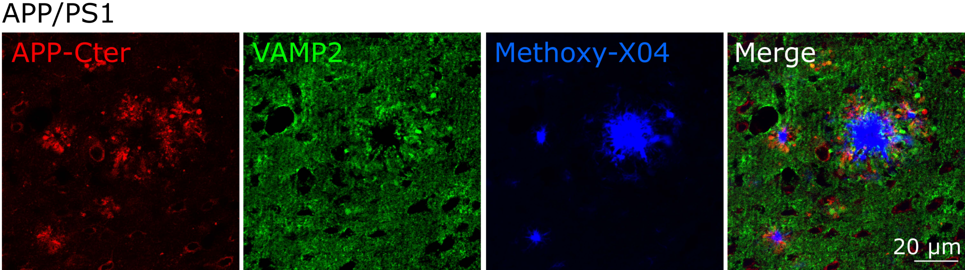
